# Gli3 utilizes Hand2 to synergistically regulate tissue-specific transcriptional networks

**DOI:** 10.1101/2020.03.13.990481

**Authors:** Kelsey H. Elliott, Xiaoting Chen, Joseph Salomone, Praneet Chaturvedi, Preston A. Schultz, Sai K. Balchand, Jeffrey D. Servetas, Aimée Zuniga, Rolf Zeller, Brian Gebelein, Matthew T. Weirauch, Kevin A. Peterson, Samantha A. Brugmann

## Abstract

Despite a common understanding that Gli TFs are utilized to reiterate a Hh morphogen gradient, genetic analyses suggest craniofacial development does not completely fit this paradigm. We demonstrated that rather than being driven by a Hh threshold, robust Gli3 transcriptional activity during skeletal and glossal development required interaction with the bHLH TF Hand2. Not only did genetic and expression data support a co-factorial relationship, but genomic analysis further revealed that Gli3 and Hand2 were enriched at regulatory elements for genes essential for mandibular patterning and development. Interestingly, motif analysis at sites co-occupied by Gli3 and Hand2 uncovered mandibular-specific, low-affinity, ‘divergent’ Gli binding motifs (**d**GBMs). Functional validation revealed these **d**GBMs conveyed synergistic activation of Gli targets essential for mandibular patterning and development. In summary, this work elucidates a novel, sequence-dependent mechanism for Gli transcriptional activity within the craniofacial complex that is independent of a graded Hh signal.

## INTRODUCTION

The Hedgehog (Hh) signaling pathway has been studied for decades in contexts ranging from organogenesis to disease (Nusslein-Volhard and Wieschaus 1980; Chang et al. 1994; Chiang et al. 1996; St-Jacques et al. 1999; Hebrok et al. 2000; Yao et al. 2002; Zhang et al. 2006). Transduction of the pathway in mammals relies on the activity of three glioma-associated oncogene (Gli) family members Gli1, 2, and 3, thought to be derived from duplications of a single ancestral gene similar to those found in lower chordates (Shin et al. 1999; Shimeld et al. 2007). While Gli2 and Gli3 transcription factors (TFs) function as both activators and repressors of Hh target genes (Dai et al. 1999; Sasaki et al. 1999; Bai et al. 2004; McDermott et al. 2005), genetic experiments have determined that Gli2 functions as the predominant activator of the pathway (Ding et al. 1998; Matise and Joyner 1999; Park et al. 2000), whereas Gli3 functions as the predominant repressor (Persson et al. 2002). All Gli family members contain five zinc-finger domains and numerous approaches (ChIP, SELEX and Protein Binding Microarray) have confirmed they all recognize a common consensus sequence, GACCACCC as the highest affinity site (Kinzler and Vogelstein 1990; Hallikas et al. 2006; Vokes et al. 2007; Vokes et al. 2008; Peterson et al. 2012). This shared consensus sequence suggests other factors and variables contribute to shaping tissue-specific and graded Gli-dependent transcriptional responses.

The fundamental and prevailing hypothesis explaining graded Hh signal transduction is the morphogen gradient (Wolpert 1969). In this model, the secreted morphogen (Sonic Hedgehog; Shh) emanates from a localized source and diffuses through a tissue to establish a gradient of activity. Responding cells are hypothesized to activate differential gene expression in a concentration dependent manner, which subsequently subdivides the tissue into different cell types. Over the years, there have been edits to the original morphogen gradient hypothesis including superimposition of a temporal variable (Dessaud et al. 2007; Dessaud et al. 2010; Balaskas et al. 2012b) and understanding how the heterogeneity in receiving cells can lead to diverse responses to the morphogen (Jaeger et al. 2004; Dessaud et al. 2008; Balaskas et al. 2012a). However, two highly studied tissues, the developing neural tube (NT) and limb, have provided the best support and understanding for the morphogen gradient as the primary mechanism used by the Hh pathway to pattern tissues.

In the NT, Shh is produced from the ventral floor plate and forms a concentration gradient along the dorsal-ventral (DV) axis that is subsequently translated into a Gli activity gradient with Gli activator (GliA) levels higher ventrally and Gli repressor (GliR) levels higher dorsally (Echelard et al. 1993; Roelink et al. 1994; Briscoe and Ericson 2001; Wijgerde et al. 2002). These opposing GliA and GliR gradients correlate with *Gli2* and *Gli3* expression patterns, respectively, and are required for patterning motor neurons and interneurons along the DV axis of the NT (Lei et al. 2004). While the most ventral cell types are lost in Gli2 mutants, Gli3 mutants have only a moderate phenotype (Ding et al. 1998; Persson et al. 2002). These observations suggest that cell identity within the ventral NT is more sensitive to levels of GliA than GliR.

In contrast, the developing limb utilizes Gli3R to perform the major patterning role, with Gli2 playing only a minor role (Hui and Joyner 1993; Mo et al. 1997; Bowers et al. 2012). Shh and Gli3R form opposing gradients across the anterior-posterior (AP) axis of the limb bud. Loss of *Gli3* results in polydactyly and a partial loss of the AP pattern, suggesting that a Gli3R gradient is necessary to determine digit number and regulate digit polarity (Wang et al. 2000; Litingtung et al. 2002; te Welscher et al. 2002). Furthermore, Gli3 is epistatic to Shh: the *Shh^-/-^;Gli3^-/-^* compound knockout has a polydactylous limb phenotype identical to the *Gli3* mutant alone, indicating that the major role of Shh in the autopod is to modulate Gli3R formation (Litingtung et al. 2002; te Welscher et al. 2002). Thus, these classic studies established the understanding that the formation of distinct Gli2 (activator) and Gli3 (repressor) gradients are necessary for converting the Hh signal transduction cascade into downstream gene expression responses within the vertebrate NT and limb.

The developing craniofacial complex represents another organ system heavily reliant upon Shh signal transduction for proper development and patterning (Helms et al. 1997; Marcucio et al. 2001; Hu et al. 2003; Cordero et al. 2004; Lan and Jiang 2009; Young et al. 2010; Xu et al. 2019); however, the mechanisms by which the craniofacial complex translates a Shh signal remain much more nebulous than those in the NT or limb. Several issues contribute to the lack of clarity in the developing face. First, rather than the simple morphology of a tube or a paddle, the facial prominences have complex morphologies that rapidly and significantly change throughout development. Second, unlike the NT and limb, patterns of *Gli2* and *Gli3* expression are not spatially distinct within the facial prominences (Hui et al. 1994). For example, despite an epithelial source of Shh on the oral axis in the developing mandibular prominence (MNP), opposing gradients of *Gli2* and *Gli3* have not been reported. Finally, conditional loss of either *Gli2* or *Gli3* alone in the neural crest cell (NCC)-derived facial mesenchyme does not result in significant patterning defects indicative of a gain-or loss-of-Hedgehog function (Chang et al. 2016). Together, these data suggest that additional mechanisms of Gli-mediated Hh signal transduction are utilized during facial development to initiate proper patterning and growth.

In this study we combined expression, genetic, genomic and bioinformatic studies to identify a novel, Gli-driven mechanism of activating tissue-specific transcriptional networks to confer positional information independent of the Hh morphogen. Specifically, Gli3 and Hand2 utilize low-affinity, divergent GBM (**d**GBM) and E-boxes to promote synergistic activation of MNP targets, outside the highest threshold of Hh signaling. We uncovered novel genetic and physical interactions between Gli3 and the bHLH TF Hand2 required within the developing MNP. Genomic binding analyses highlighted enrichment of both factors at the same CRMs and revealed a surprising, motif-dependent synergism distinct to Gli3 and Hand2. Importantly, this synergism is required for robust activation of Gli targets important for mandibular patterning, glossal development and skeletogenesis. Our findings suggest that context-dependent optimization of Gli binding site occupancy in the presence of Hand2 is critical for modulating tissue-specific transcriptional output within a tissue that lacks an obvious Shh morphogen gradient. Hence, these findings define how craniofacial prominences can serve as distinct developmental fields that interpret Hh signals in a manner unique to other organ systems.

## RESULTS

### Loss of Gli TFs and Hand2 *in vivo* generates micrognathia and aglossia

To attain a comprehensive understanding of Gli TF function during craniofacial development, we generated conditional mutant mice lacking *Gli2* and *Gli3* in the NCC-derived mesenchyme (*Gli2^f/f^;Gli3^f/f^;Wnt1-Cre,* herein referred to as *Gli2/3* cKO). During our analysis we observed a severe micrognathia phenotype in *Gli2/3* cKO embryos that was highly reminiscent of those previously described for *Hand2^f/f^;Wnt1-Cre* (*Hand2* cKO) mutants (Morikawa et al. 2007; Barron et al. 2011). Relative to wild-type embryos, *Gli2/3* cKO mutants presented with low-set pinnae, aglossia and micrognathia (Fig. 1A-C’, I). Posterior cranial skeletal structures including the tympanic ring, hyoid bone, coronoid, condylar, and angular processes were hypoplastic, and incisors were absent (Figure 1D, I, Supp. Fig. S1A-B). Interestingly, conditional loss of either *Gli2* or *Gli3* alone (*Gli2^f/f^;Wnt1-Cre* or *Gli3^f/f^;Wnt1-Cre*) did not replicate the mandibular phenotype observed in double mutants (Chang et al. 2016).

**Figure 1.**
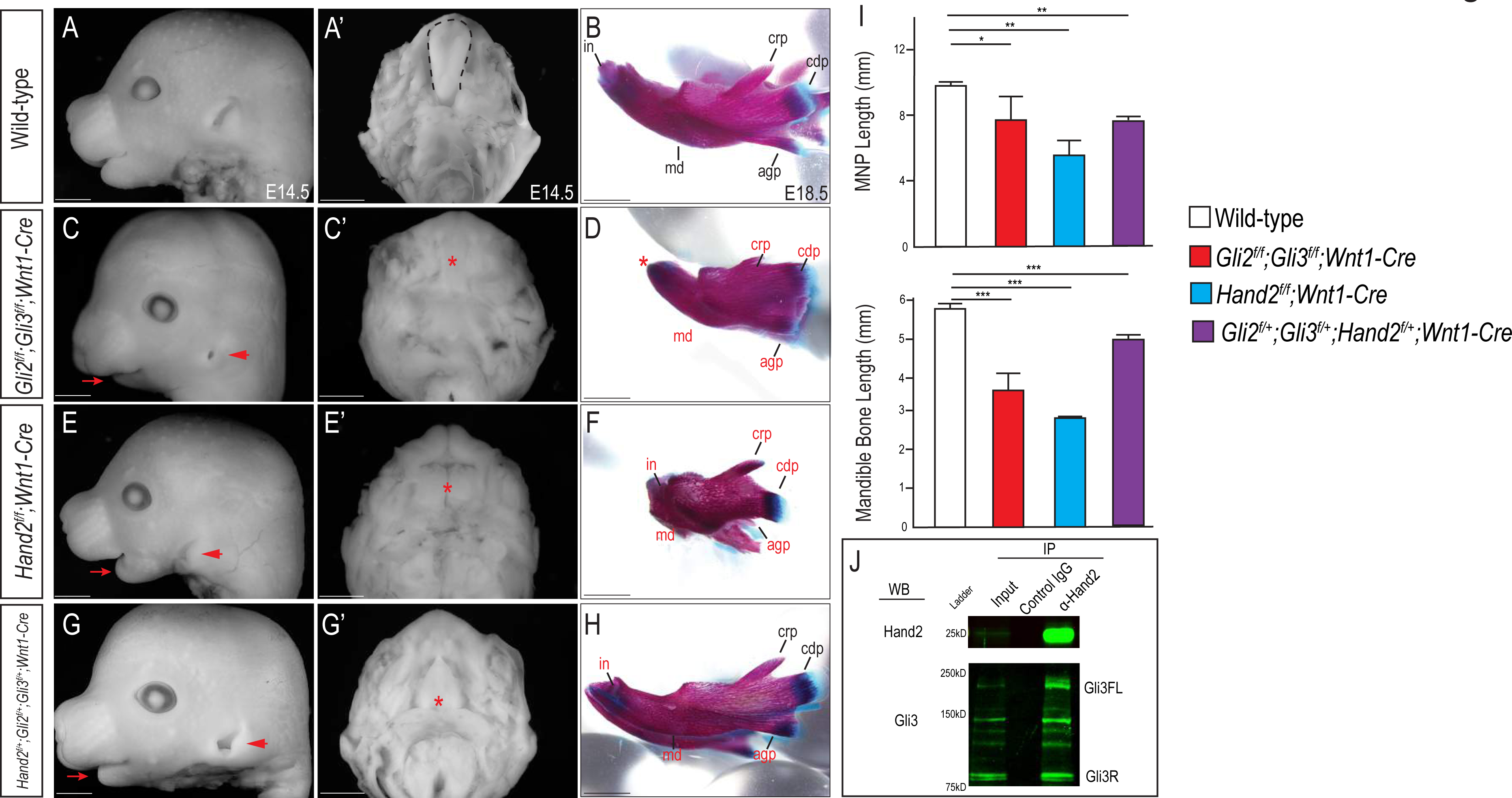
Gli and Hand2 are required for mandibular developmental networks in vivo (A,C,E,G) Lateral cranial view or (A’,C’,E’,G’) dorsal mandibular view of wild-type, *Gli2^f/f^;Gli3^f/f^;Wnt1-Cre*, *Hand2^f/f^;Wnt1-Cre*, and *Gli2^f/+^;Gli3^f/+^;Hand2^f/+^;Wnt1-Cre* embryos at E14.5. Red arrow indicates micrognathia. Red arrowhead indicates low-set pinnae. Dotted black line denotes tongue and red asterisk highlights observed aglossia. (B,D,F,H) Lateral cranial view of Alizarin Red and Alcian Blue staining to mark bone and cartilage respectively in wild-type, *Gli2^f/f^;Gli3^f/f^;Wnt1-Cre*, *Hand2^f/f^;Wnt1-Cre*, and *Gli2^f/+^;Gli3^f/+^;Hand2^f/+^;Wnt1-Cre* mandibles at E18.5. Abbreviations: md, mandible; in, incisor; crp, coronoid process; cdp, condylar process; (I) Measurements of MNP and mandibular bone. Data are expressed as mean +SD. *p<0.05, **p<0.01, ***p<0.001. (J) Co-immunoprecipitation showing interaction between Gli3 and Hand2 within E10.5 MNPs. Scale bar: 1mm. See also Supplemental Figure 1.

Similar to *Gli2/3* cKO embryos, and consistent with previous reports, we observed low-set pinnae, micrognathia and aglossia in *Hand2* cKO mutants (Barron et al. 2011) (Fig. 1E-E’, I). Skeletal analysis of *Hand2* cKO mutants confirmed a dysmorphic and hypoplastic mandible and loss of Meckel’s cartilage (Fig. 1F’, I), similar to the *Gli2/3* cKO mutant phenotype. Concurrent with *Gli2/3* cKO mutants, *Hand2* cKO mutants had dysmorphic mandibles, and posterior structures such as the tympanic ring and angular processes were lost, while the coronoid and condylar processes were hypoplastic (Supp. Fig. S1C). Although the hyoid bone was present, it was abnormally fused to middle ear cartilage and underwent excessive/ectopic ossification (Barron et al. 2011).

To determine if the phenotypic similarities between *Gli2/3* cKO and *Hand2* cKO mutants were due to an epistatic relationship between Gli TFs and Hand2, we analyzed gene expression in mutant embryos by RNA-seq. We did not detect significant changes in expression in *Hand2* or any Shh pathway components in *Gli2/3* cKO and *Hand2* cKO mutant MNPs, respectively (Supp. Table 1). Therefore, the similarities of the mandibular phenotype along with the maintained expression of each TF in each respective mutant, suggested that these TFs may work in parallel to promote patterning and development of the MNP.

To test the hypothesis that Gli TFs and Hand2 regulate a common transcriptional network within NCCs of the MNP, we performed combinatorial genetic and biochemical experiments. First, while heterozygous *Gli2/3* or *Hand2* conditional mutants (*Gli2^f/+^;Gli3^f/+^;Wnt1-Cre* or *Hand2^f/+^;Wnt1-Cre,* respectively) did not produce severe MNP phenotypes (Supp. Fig. S1D-E’), triple heterozygotes (*Hand2^f/+^;Gli2^f/+^;Gli3^f/+^;Wnt1-Cre*) resulted in MNP phenotypes similar to those observed in the full *Gli2/3* cKO or *Hand2* cKOs, including low-set pinnae, micrognathia, smaller incisors, and aglossia (Fig. 1G-H’, I). Thus, these genetic experiments supported the possibility that Gli TFs and Hand2 function together to direct MNP development. Second, to determine if Hand2 and Gli TFs physically interact *in vivo*, we performed co-immunoprecipitation using embryonic day (E) E10.5 wild-type MNPs. Hand2 physically interacted with both full-length and truncated isoforms of Gli3, but only the truncated isoform of Gli2 (Fig. 1J, Supp. Fig. S1F). Taken together, these data provided genetic, molecular and biochemical evidence suggesting that Gli and Hand2 TFs participate within a common transcriptional network important for mandibular development, and further suggest that there may be a unique role for Gli/Hand2 cooperation.

### Gli2, Gli3, and Hand2 are co-expressed NCC-derived populations that give rise to skeletal and glossal progenitors

To explore the molecular basis for Gli-mediated micrognathia and investigate the hypothesis that Gli TFs and Hand2 cooperate to initiate MNP patterning and development, we examined the endogenous expression of these TFs during early MNP development using single molecule fluorescent *in situ hybridization* (RNAscope). Contrary to the distinct and opposing *Gli*2 and *Gli3* expression domains observed in other developing organ systems (Lee et al. 1997; Sasaki et al. 1997; Buscher and Ruther 1998; Lei et al. 2004), no spatial distinction or opposing expression gradients were observed between *Gli2* and *Gli3* in the developing MNP (Fig. 2A-C’). Furthermore, *Gli2* and *Gli3* were co-expressed within many cells of the developing MNP (Fig. 2C-C’), supporting the hypothesis that the developing MNP uses unique mechanisms to integrate spatiotemporal information.

**Figure 2.**
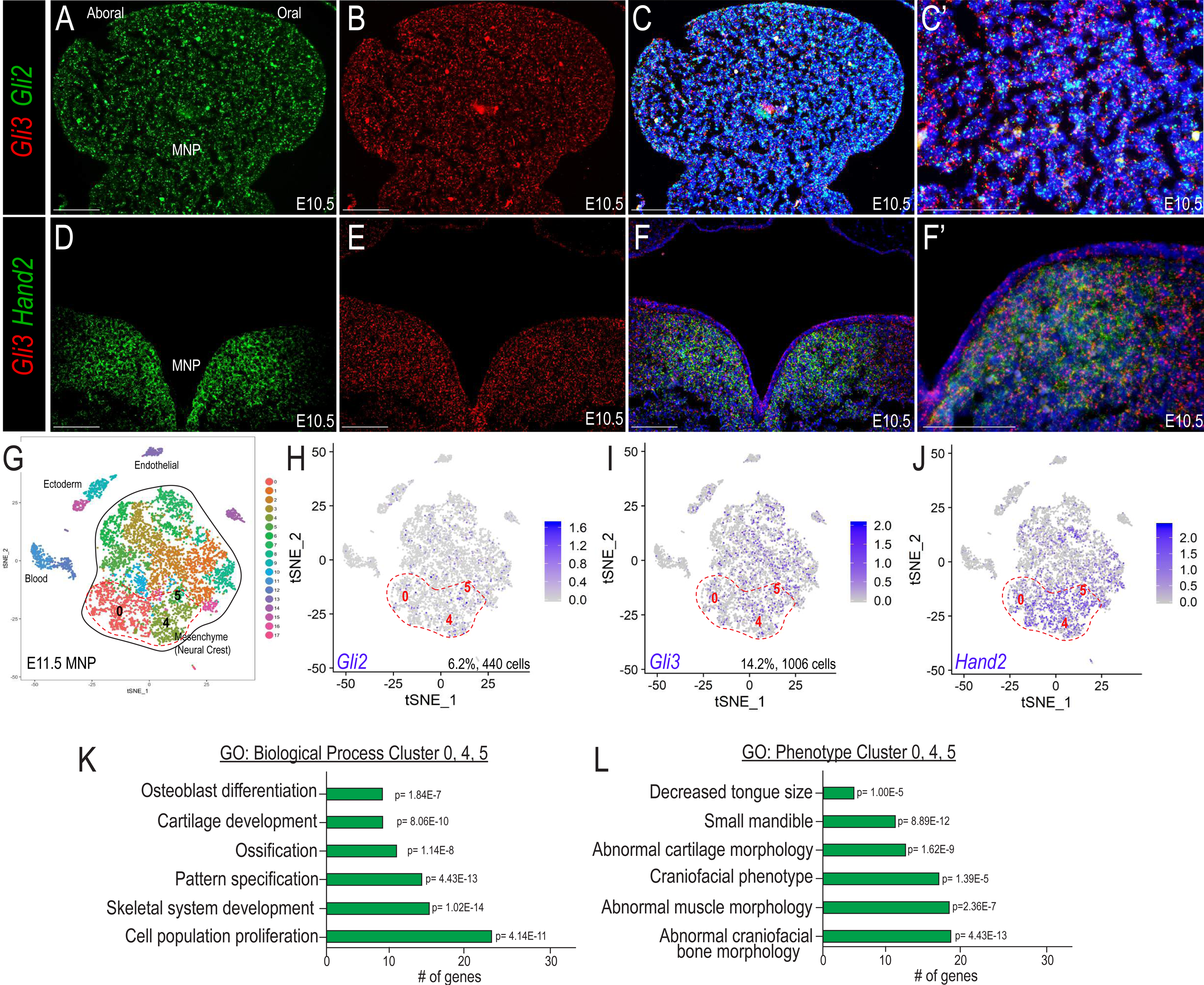
Gli expression in the developing MNP suggests other inputs are required to coordinate diversity of outputs (A-C) Expression of Gli2 and Gli3 within the developing MNP as revealed by smFISH on sagittal sections of E10.5 embryos. (C’) Higher magnification of C. (D-F) Expression of Gli3 and Hand2 within the developing MNP as revealed by smFISH on frontal sections of E10.5 embryos. (F’) Higher magnification of F. (G) tSNE plot of single cell RNA-sequencing of the E11.5 MNP. (H-J) Single cell expression of Gli2, Gli3, and Hand2 in the E11.5 MNP. Dotted red line indicates Gli+/Hand2+ NCC clusters (0, 4, 5). (K-L) GO terms associated with marker genes for clusters 0, 4, 5 indicate Gli+/Hand2+ cells contribute to skeletogenesis and glossal development. See also Supplemental Figure 2.

As opposed to the widespread MNP expression of *Gli3* and *Gli2, Hand2* expression was confined to the medial aspect of the MNP (Fig. 2D-E) (Srivastava et al. 1997; Thomas et al. 1998; Barron et al. 2011; Funato et al. 2016). Interestingly, while many *Gli3*+ cells did not express *Hand2*, most or all *Hand2*+ cells co-express *Gli3* (Fig. 2F, F’). To determine the identity of cells co-expressing *Gli2/3* and *Hand2*, we performed single-cell RNA-sequencing in the developing MNP. At E11.5, unsupervised clustering identified 17 distinct clusters in the MNP, including a central grouping of mesenchymal clusters derived from NCCs (Fig. 2G; Supp. Fig. S2A-C). Coincident with RNAscope, *Gli2* and *Gli3* expression were not restricted to, nor enriched in any particular cell cluster. While we failed to observe a gradient or polarized expression of Gli TFs throughout the MNP, there were over 2-fold more cells expressing *Gli3* compared to *Gli2* (Fig. 2H-I, 1006 cells, 14.2% versus, 440 cells, 6.2%). Thus, despite earlier studies suggesting that the MNP is patterned by Gli activation traditionally hypothesized to occur via Gli2 (Jeong et al. 2004; Millington et al. 2017), these data suggested a more extensive role for Gli3. Contrary to the widespread *Gli* TF expression*, Hand2* was enriched in NCC-derived clusters 0, 4, and 5 (Fig. 2J). GO enrichment analyses for clusters 0, 4, and 5 revealed that these NCC-derived cells contributed to processes altered in *Gli2/3* cKO and *Hand2* cKO mutant embryos, such as skeletal and glossal development, and pattern specification (Fig. 2K). Additionally, phenotypes arising from dysregulation of these cell clusters included decreased tongue size and small mandible (Fig. 2L), suggesting these *Gli/Hand2* co-expressing clusters may be responsible for phenotypes present in conditional knockouts (Fig. 1). Since a *Gli2/3* expression gradient or restriction from cell types cannot explain diverse Gli-dependent transcriptional outputs, we hypothesized that functional interactions with Hand2, which is required for development of similar MNP tissues (Fig. 1), may explain this phenomenon.

### Gli3 and Hand2 co-regulate targets in the developing MNP

To determine if Gli TFs and Hand2 regulated a common group of target genes, we performed bulk RNA-sequencing on E10.5 *Gli2/3* cKO and *Hand2* cKO MNPs (GSE141431). Transcriptome profiling and GO analyses revealed a wide variety of differentially expressed genes affecting a number of biological processes including muscle system process, anterior/posterior patterning, regionalization, and cell-cell signaling (Fig. 3A-B). These enriched GO terms reflected the phenotypes (e.g., aglossia, micrognathia) in *Gli2/3* cKO or *Hand2* cKO mutant embryos and were consistent with the *Gli+/Hand2+* cell types identified by scRNA-seq. Furthermore, hypergeometric tests revealed enrichment of shared transcripts, with 50% of genes differentially expressed in *Gli2/3* cKO MNPs also differentially expressed in *Hand2* cKO MNPs (Fig. 3C, p=3.7E-284). This highly significant overlap led us to further investigate mechanisms of a possible co-factorial relationship between Gli TFs and Hand2.

**Figure 3.**
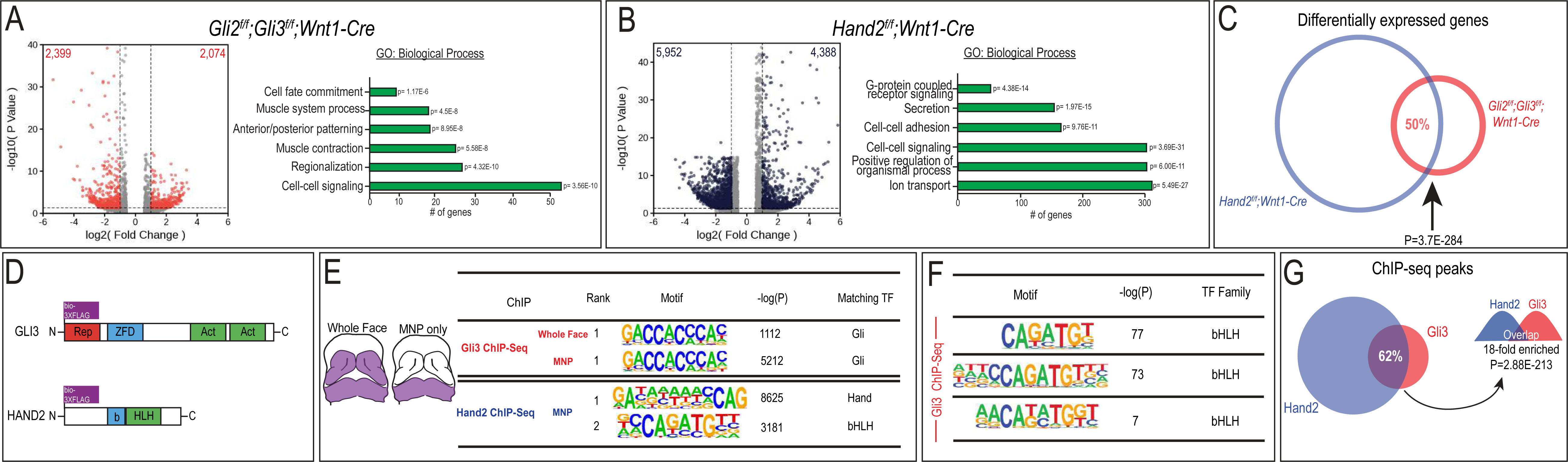
Gli3 and Hand2 regulate and bind to a common set of target genes in the MNP (A-B) Volcano plots and GO terms associated with differentially expressed genes from *Gli2^f/f^;Gli3^f/f^;Wnt1-Cre* or *Hand2^f/f^;Wnt1-Cre* E10.5 MNPs. Number of genes with significant differential expression (fold change > 1.5, adjusted p-value < 0.05) indicated on each panel. (C) Venn diagram of shared differentially expressed genes in *Gli2^f/f^;Gli3^f/f^;Wnt1-Cre* and *Hand2^f/f^;Wnt1-Cre* MNPs. (D) Endogenously FLAG-tagged mice used for in vivo ChIP-seq.

Next, we assessed whether Gli TFs and Hand2 occupied the same CRMs by performing ChIP-seq analyses *in vivo* using endogenously FLAG-tagged alleles for each TF (Lopez-Rios et al. 2014; Osterwalder et al. 2014; Lorberbaum et al. 2016) (Fig. 3D). Since our previous biochemical and expression data supported a unique role for Gli3 in the MNP and a unique relationship between Gli3 and Hand2, we focused our characterization of genomic occupancy on Gli3. As expected, the most highly enriched TF binding site observed in Gli3 ChIP-seq on either E11.5 whole face (frontonasal, maxillary and mandibular prominence) or MNPs alone reflected the previously reported ‘canonical’ Gli binding motif (**c**GBM) defined by the GACCACCC 8-mer (Kinzler and Vogelstein 1990; Vokes et al. 2008) (Fig. 3E). Similarly, Hand2 peaks contained both canonical bHLH E-box motifs (CANNTG) and Hand-specific E-box motifs (Maves et al. 2009; Kulakovskiy et al. 2013) (Fig. 3E). Further motif enrichment analyses revealed that bHLH motifs were also significantly enriched within Gli3 MNP peaks (Fig. 3F). Comparison between Gli3 and Hand2 MNP ChIP-seq peaks via regulatory element locus intersection (RELI) (Harley et al. 2018) revealed a significant overlap of genomic locations occupied by Gli3 and Hand2 in the MNP (Fig. 3G, 62%, 18-fold enriched, p=2.88E-213). These data provided evidence that Gli3 and Hand2 were enriched at similar CRMs within common transcriptional networks.

To determine the biological relevance of combined Gli3/Hand2 transcriptional input during MNP development, we performed scRNA-seq on MNPs at E13.5, a stage when NCC differentiation into distinct cell types has begun (Fig. 4A) (GSE141173). Similar to E11.5 analyses, *Gli3* expressing cells (16.8%) were more prevalent than *Gli2* expressing cells (10.4%), and observed in almost all clusters of the E13.5 MNP (Fig. 4A-B, Supp. Fig. S3A), and *Hand2* expressing cells were restricted to a subset of NCC-derived mesenchymal clusters (Fig. 4C). Co-expression of *Gli3* and *Hand2* was observed within osteogenic (cluster 5) and glossal musculature (clusters 1, 15, 19) clusters. Integration analysis of our E11.5 and E13.5 MNP scRNA-seq data revealed that the most significantly impacted cell states in *Gli2/3* cKO or *Hand2* cKO (NCC-derived skeletal elements and glossal muscle) mutants originated from cells that co-expressed *Gli3* and *Hand2* at E11.5 (E11.5 clusters <u>0, 4, 5</u>; E13.5 clusters 1,5,15; Fig. 3H, Supp. Fig. S3B-C). These data further supported the hypothesis that Gli3/Hand2 transcriptional input was an essential driver for proper MNP patterning and development.

**Figure 4.**
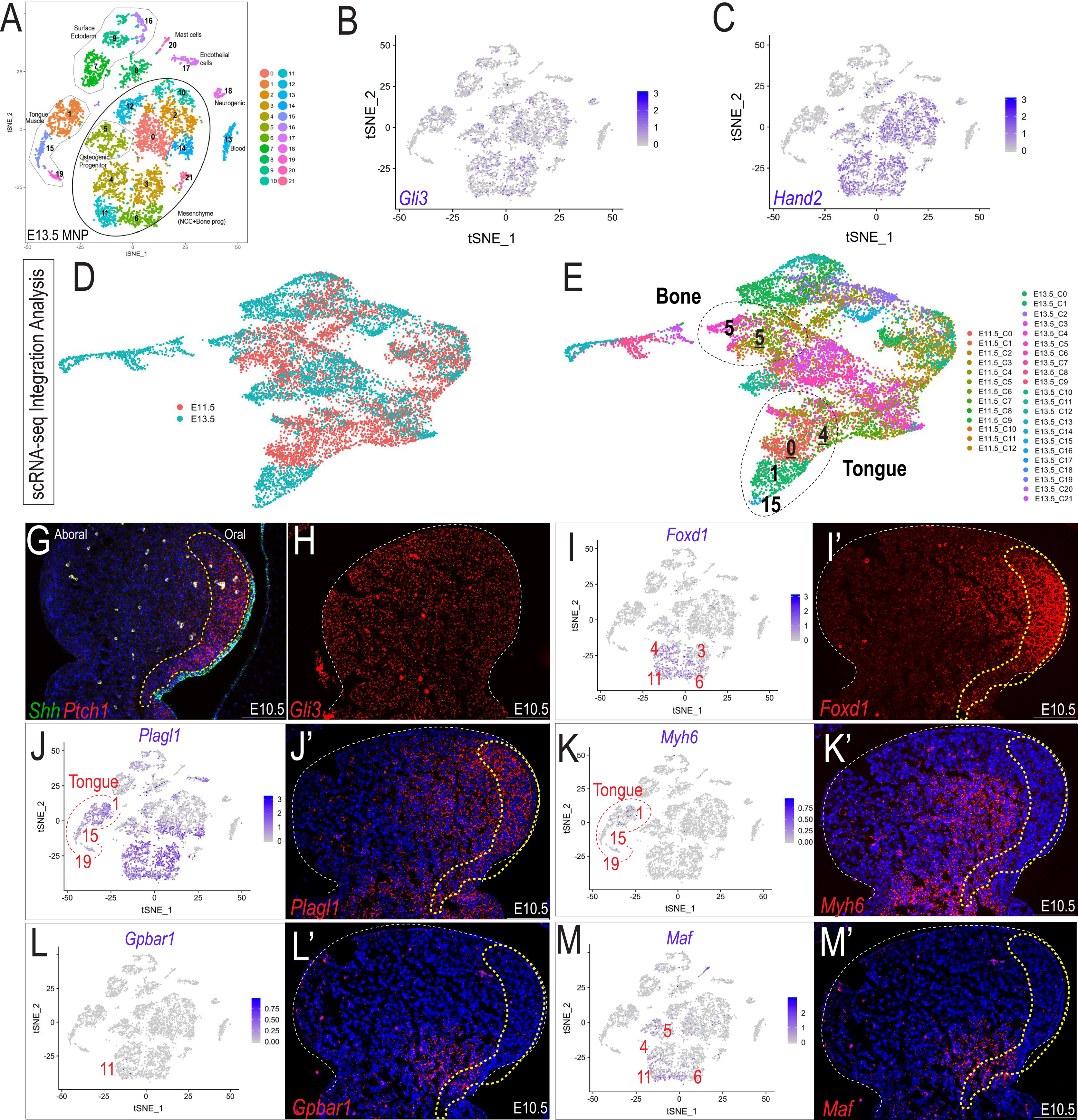
Hand2 and Gli3 coordinate expression of muscular and skeletal gene regulatory networks (A) tSNE plot of single cell RNA-sequencing from E13.5 wild-type MNP. (B-C) Single cell expression of *Gli3* and *Hand2* in E13.5 MNP. (D-E) UMAP plot after integration analysis and re-clustering of E11.5 and E13.5 scRNA-seq MNP samples showed the E13.5 muscle (1,15) and osteogenic (5) clusters are most similar to, and likely derived from E11.5 *Gli3+/Hand2+* NCC clusters (0,4,5). (G-H) Expression of *Shh, Ptch1*, and *Gli3* as revealed by smFISH in sagittal sections of E10.5 MNPs. Dotted yellow line indicates highest Shh-responsive area marked by *Ptch1*. (I-M’) scRNA-seq and smFISH expression of *Gli3* and *Hand2* targets involved with MNP patterning. See also Supplemental Figure 3.

To confirm that Gli3/Hand2 transcriptional input was necessary for MNP patterning, skeletogenesis and muscular development/glossogenesis, we selected 5 target genes identified in our transcriptome and ChIP-seq analyses and examined their expression patterns, relative to *Shh* and *Ptch1* expression. *Forkhead Box d1* (*Foxd1*), a well-characterized Gli target in the MNP involved in regionalization (Jeong et al. 2004) was enriched in the *Gli3+/Hand2+* clusters 3, 4, 6, 11 and expressed both within and outside the *Ptch1* domain in NCC-derived mesenchyme (Fig. 4G-I’). *Pleiomorphic adenoma gene-like 1* (*Plagl1*), a gene which impacts glossal development (Li et al. 2014) and *Myosin heavy chain 6* (*Myh6*), a myosin isoform found in specialized skeletal muscles (Lee et al. 2019) were expressed within *Gli3+/Hand2+* descendent tongue/musculature clusters 1, 15, and 19 (Fig. 4J-K’,), and both within and outside the *Ptch1* expression domain. Finally, *G-protein coupled bile acid receptor 1* (*Gpbar1*, a.k.a. *Tgr5*), a gene involved in osteoblast differentiation and mineralization (Wang et al. 2018) and *Avian Musculoaponeurotic fibrosarcoma oncogene homolog* (*Maf*), a TF involved in chondrocyte differentiation (Hong et al. 2011), were expressed in *Gli3+/Hand2+* clusters 4, 5, 6, and 11 corresponding with NCC-derived mesenchyme and skeletal progenitors (Fig. 4L-M’), and both within and outside the *Ptch1* expression domain. Thus, transcriptome (Fig. 3A-C), genome occupancy, (Fig. 3D-G) and gene expression (Fig. 4) data all suggested that Hand2 and Gli3 collaborate at common CRMs to activate common networks within the MNP to establish osteogenic, chondrogenic, and glossal/muscle cell fates. These data further suggested that despite being Gli3 targets, these genes do not require graded Shh activity for expression, but rather are primarily reliant upon combined input of Gli3/Hand2.

### Low-affinity Gli binding motifs are within close proximity to E-boxes and specific to the developing mandible

To examine mechanisms of transcriptional regulation for shared Gli3/Hand2 targets, we performed *de novo* motif analysis on Gli3-alone versus Gli3/Hand2-overlaping peak regions. As expected, the most enriched motif within Gli3-alone peaks was the previously reported ‘canonical’ GBM (**c**GBM) defined by the ‘GACCACCC’ 8-mer (Kinzler and Vogelstein 1990), which was 9.8-fold-enriched compared to background sequences (Fig. 5A). Surprisingly, when we performed motif analysis on overlapping peaks shared between Gli3 whole face and Hand2 MNP samples, the top-ranked GBM (6.3-fold-enriched over background sequences) deviated from the **c**GBM 8-mer, with the most notable change being the reduced weight of the highly conserved ‘A’ at the 5^th^ position (Fig. 5A’). To specifically address the Gli3/Hand2 relationship in the MNP, we repeated these analyses using only overlapping peaks from Gli3 and Hand2 MNP samples. Here, the top-ranked GBM present (6.2-fold-enriched over background) differed even further from the canonical 8-mer, having a close to equal probability of either a ‘T’ or an ‘A’ at the highly constrained 5^th^ position (Fig. 5A’’). We designated this GACC**<u>T</u>**CCC 8-mer as a ‘divergent’ GBM (**d**GBM). Interestingly, the **d**GBM was most clearly revealed upon comparisons of MNP data sets. These data supported the possibility that the **d**GBM utilized by Gli3 and Hand2 was specific to the MNP. To test this hypothesis, we repeated our *de novo* motif analysis comparing to publicly available data from the developing limb (Fig. 5B; GSE55707) (Osterwalder et al. 2014). Strikingly, the **d**GBM present in our MNP analysis was not present when comparing Hand2 binding in the limb. Rather, these analyses revealead the highly constrained 5^th^ position remained exclusively a heavily weighted ‘A’. Together, these data suggested a tissue-specific role for this MNP-enriched **d**GBM.

**Figure 5.**
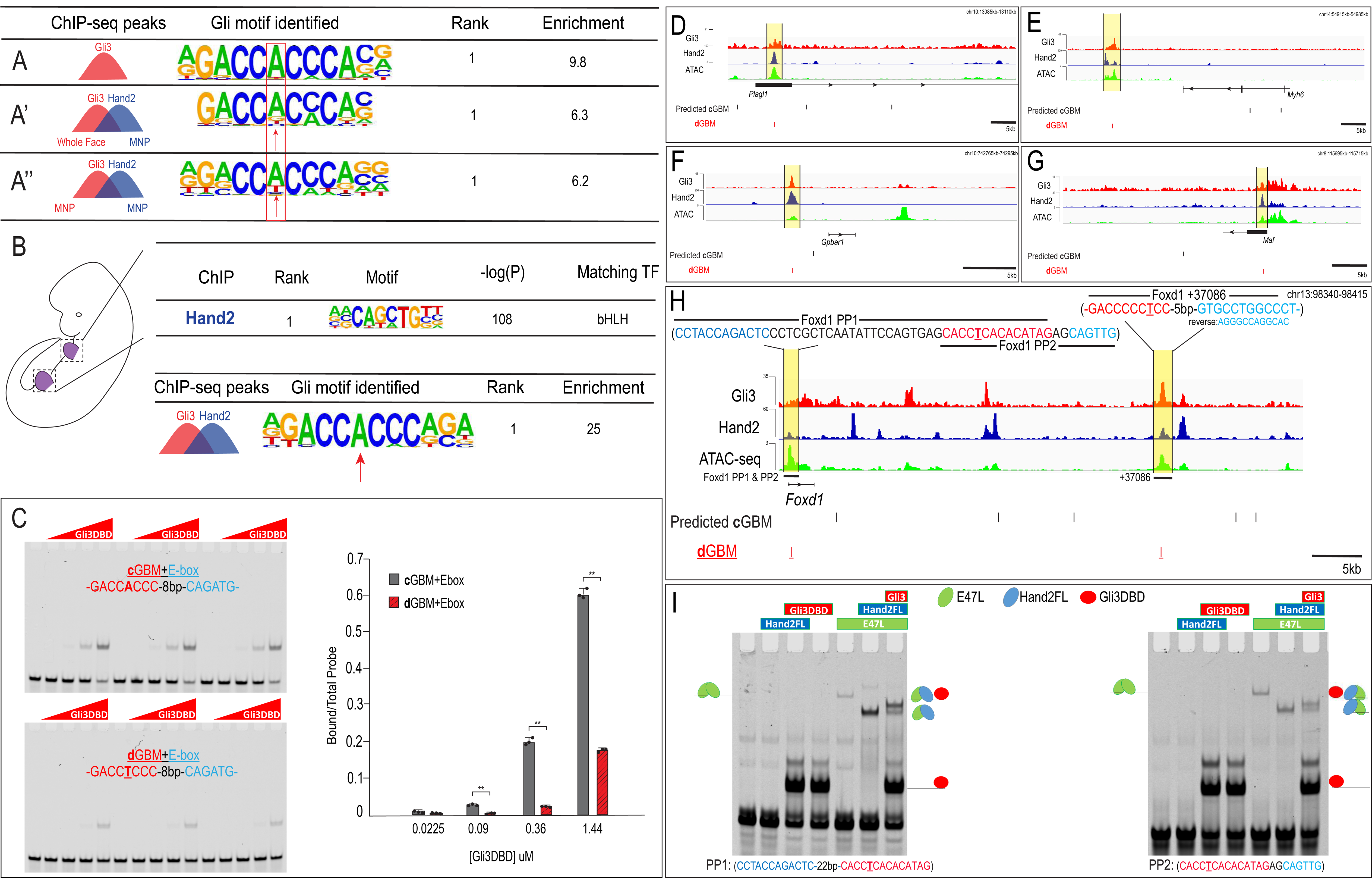
Low affinity divergent Gli binding motifs are found near E-boxes (A-A’’) De novo motif enrichment for Gli3-only peaks in MNP, Gli3/Hand2 overlapping peaks, comparing (A’) Gli3-whole face peaks to Hand2 MNP peaks or (A’’) Gli3 MNP peaks to Hand2 MNP peaks. (B) (Top) Known motif enrichment of Hand2 peaks from limb buds of endogenously FLAG-tagged mice. (Bottom) De novo motif enrichment of Gli3/Hand2 overlapping peaks, comparing Gli3 peaks from whole face and Hand2 peaks from limb. (C) Electrophoretic mobility shift assay (EMSA) and quantification of affinity showing that the Gli3 DNA binding domain (Gli3DBD) binds with increased affinity to canonical GBMs (**c**GBM) relative to divergent GBMs (**d**GBMs). Results used for quantification are shown in triplicate. (D-H) Overview of MNP-specific regulatory input to the *Plagl1, Myh6, Gpbar1, Maf* and *Foxd1* locus. **c**GBMs (black line) and **d**GBMs (red lines) are indicated below the signal tracks for Gli3 (red) and Hand2 (blue) ChIP-seq and ATAC-seq (green). PP1=promoter proximal 1, PP2=promoter proximal 2. (I) EMSA for PP1 and PP2 using Gli3DBD, Hand2 full-length (FL), and E47L/Tcf3, the Hand2 heterodimerization partner. Gli3DBD bound these regulatory regions alone while Hand2 binds in the presence of heterodimerization partner E47L. Binding of all three factors resulted in a supershift of the complex. The observed complexes are illustrated for clarity. See also, Supplemental Figure 4.

To confirm the decreased frequency of **c**GBM binding events in the presence of Hand2 in the MNP, we quantified the incidence of the **c**GBM 8-mers using a strict counting method. While the consensus **c**GBM 8-mer (GACCACCC) was detected in 16% of Gli3-only peaks collected from the MNP, its occurrence was significantly reduced to only 2% of Gli3/Hand2 overlapping peaks. Additionally, while the **c**GBM was the 33^rd^ most frequent 8-mer in Gli3-only MNP peaks (out of 32,896 possibilities), it was 563^rd^ in frequency in Gli3/Hand2 overlapping MNP peaks (Supp. File 1). This finding, in conjunction with the motif enrichment results, further supported a deviation from the **c**GBM when Hand2 and Gli3 peaks overlapped.

To examine the effect an ‘A’ to ‘T’ transition at the 5^th^ position had on relative binding affinity, we utilized previously published Gli3 protein binding microarray (PBM) E-score data (Peterson et al. 2012). PBM E-scores range from -0.5 to +0.5, with values above 0.4 generally considered strong binding sites (Berger et al. 2006; Berger and Bulyk 2009). Interestingly, substitution of ‘A’ to ‘T’ in the 5^th^ position of comparable 8-mers reduced the E-score for Gli3 binding from 0.42 to 0.33, indicating that Gli3 alone would have a lower affinity for the **d**GBM sequence. Likewise, previous studies in *Drosophila* reported that low-affinity non-canonical GBMs with a ‘T’ in the 5^th^ position, similar to what we term the **d**GBM, were responsible for regulating broad expression of Ci targets in zones of lower Hh signaling (Parker et al. 2011). To directly test the binding affinity of Gli3 to a **d**GBM, we performed gel shift assays on synthetic sequences containing either a **d**GBM+E-box or **c**GBM+E-box. These experiments confirmed that a single nucleotide alteration from ‘A’ to ‘T’ in the 5^th^ position significantly decreased the affinity of Gli3 DNA binding (Fig. 5C). Together, these data confirmed the identification of distinct, low affinity **d**GBMs enriched at genomic loci bound by both Gli3 and Hand2 within the MNP.

Next, we set out to confirm the presence and utility of endogenous **c**GBMs and **d**GBMs within Gli3 targets in the MNP. Using the Cis-BP database (Weirauch et al. 2014), we predicted high-affinity **c**GBMs either upstream or within previously identified Gli3/Hand2 targets, including *Plagl1, Myh6, Gpbar1, Maf,* and *Foxd1*. To identify regions with regulatory potential, we integrated these Cis-BP-identified **c**GBMs with our MNP-specific ATAC-seq and ChIP-seq data to highlight regions of open chromatin that were bound by Gli3 and Hand2. Interestingly, we frequently saw areas of open chromatin occupied by Gli3 and Hand2 that did not contain a high-affinity **c**GBM (Fig. 5D-H). This was in stark contrast to the Gli3-bound areas of open chromatin heavily populated with **c**GBMs at the *Ptch1* locus (Supp. Fig. S4A, black lines). Instead, these loci all displayed Gli3 and Hand2-bound regions containing **d**GBMs (Fig. 5D-H, red lines).

Analysis of the *Foxd1* regulatory landscape revealed a potential regulatory region near the *Foxd1* promoter that contained a **d**GBM 22 base pairs downstream of an E-box and 2 base pairs upstream of a second E-box (Fig. 5H). We tested each combination of this region (**d**GBM + downstream E-box or **d**GBM + upstream E-box), designating them as promoter proximal 1 (PP1) and promoter proximal 2 (PP2). The second putative regulatory region was downstream of the *Foxd1* coding region at +37086. Due to varying observable orientations and spacings between GBMs and E-boxes, we tested if there was a statistical preference for any single spacing or orientation of GBMs and E-boxes inside of Gli3/Hand2 overlapping genomic regions using the previously published COSMO method (Narasimhan et al. 2015). Despite identifying 628 “intersecting peaks” that contain a GBM and Hand2 motif within 100 bases of each other, no particular spacing/configuration was present in greater than ∼0.2% of sequences (Supp. Fig. S4B). These findings suggested flexibility in the regulatory architecture governing the spacing and orientation of Gli3 and Hand2 binding sites within CRMs bound in the developing MNP.

While ChIP data was highly suggestive of Gli3/Hand2 co-occupancy at regulatory regions containing **d**GBM and E-box motifs, it did not test if Gli3 and Hand2 were able to simultaneously bind an endogenous **d**GBM and an adjacent E-box. To ask this question, we performed gel-shift assays with the Gli3 DNA-binding domain and full-length Hand2 (Hand2FL). Gel-shift analysis revealed that the Gli3 DNA-binding domain independently binds the **d**GBM present in PP1 and PP2 (Fig. 5I). In addition, we found that Hand2 could not independently bind the E-box motifs present in the PP1 and PP2 probes but could bind as a heterodimer in the presence of E47L (Tcf3), an E-protein bHLH that cooperatively binds DNA with many tissue-specific bHLHs (Fig. 5I). This is consistent with reports that binding of many bHLH TFs require dimerization with other widely expressed E-protein family members (Firulli 2003). Importantly, Gli3 and Hand2 were able to simultaneously bind **d**GBM/E-box regions within both PP1 and PP2 (Fig. 5I), in a dose-dependent manner (Supp. Fig. S4C). Together, these data suggested that Gli3 and Hand2 simultaneously occupy potential regulatory regions containing a low-affinity **d**GBM and an E-box. We next sought to investigate how Gli3/Hand2 cooperation impacted transcriptional output.

### Gli3 and Hand2 synergize at dGBMs

To examine Gli3/Hand2 transcriptional activity, synthetic luciferase reporter constructs containing either the **c**GBM or a **d**GBM plus an E-box (**c**GBM+E-box, **d**GBM+E-box, respectively) were transfected into the cranial NCC line, O9-1 (Ishii et al. 2012). Luciferase activity was measured after transfection of Gli3 alone, Hand2 alone or Gli3 and Hand2 together (Fig. 6A; Supp. Fig. S5A). Regardless of the GBM present, expression of either Gli3 alone or Hand2 alone significantly elevated the luciferase activity of reporter constructs relative to control conditions (Fig. 6A). Co-expression of Gli3 and Hand2 with the **c**GBM+E-box synthetic reporter resulted in a small, but significant increase in luciferase expression compared to either Gli3 or Hand2 alone. In stark contrast, co-expression of Gli3 and Hand2 in the presence of the **d**GBM+E-box synthetic reporter resulted in a significant and synergistic (more than additive) upregulation of luciferase expression compared to either Gli3 or Hand2 alone (Fig. 6A). Together, these results indicated that 1) the low-affinity **d**GBM conveyed a distinct function from the **c**GBM, 2) low-affinity the **d**GBM+E-box produced synergistic transcriptional output in the presence of Gli3 and Hand2 and 3) synergistic activity was independent of a graded Hh signal, since the response was observed without Hh stimulation.

**Figure 6.**
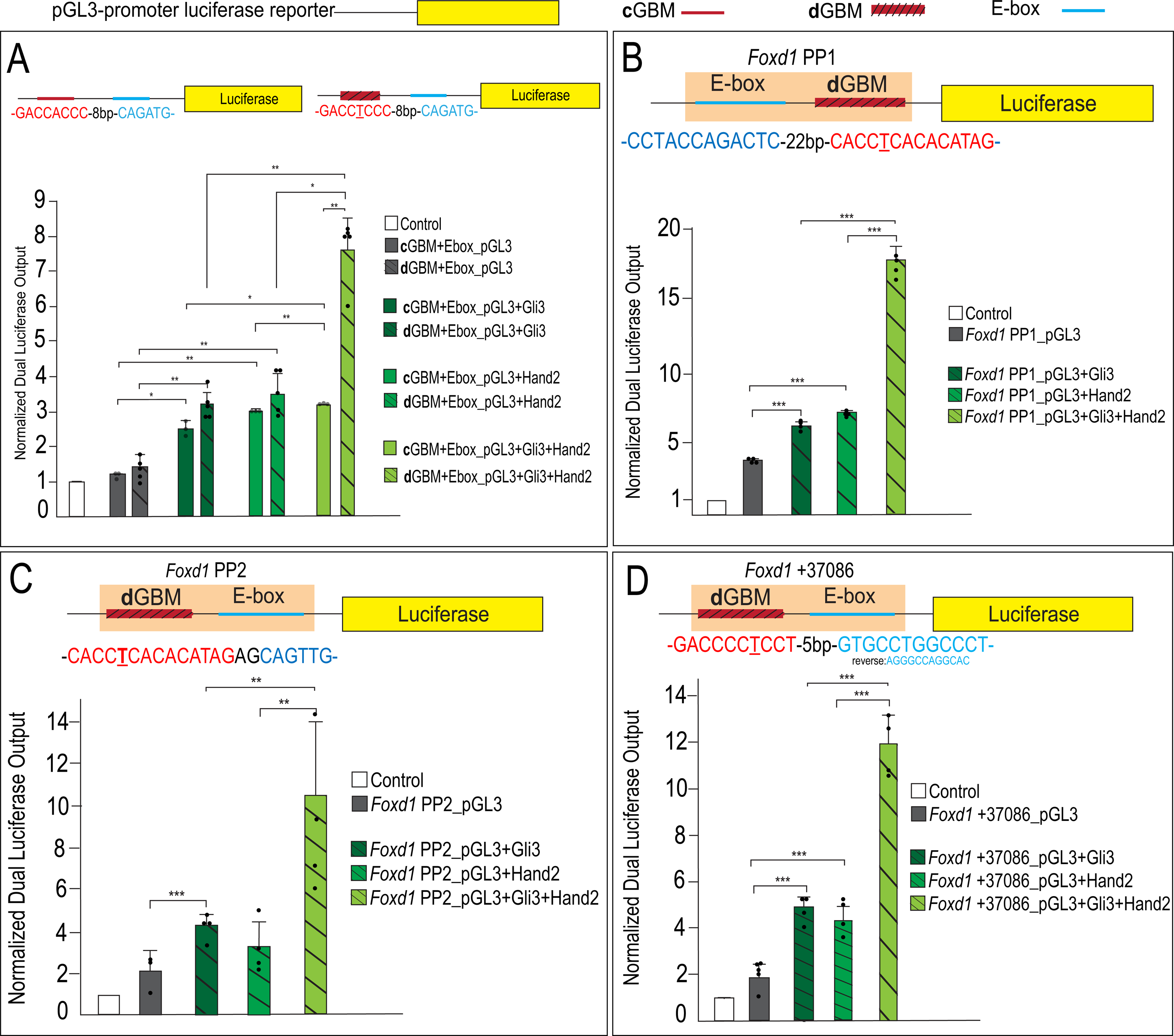
Gli3 and Hand2 synergistically activate low-affinity **d**GBMs (A) Luciferase reporter activity of synthetic constructs containing a **c**GBM and E-box (solid bars) or **d**GBM and E-box (hatched bars) in response to transfection of Gli3, Hand2, or both in O9-1 cells. (B-D) Luciferase reporter activity of the endogenous *Foxd1* putative regulatory region fragments PP1, PP2, and +37086 after transfection with Gli3, Hand2, or both in O9-1 cells. Data are expressed as mean + SD with biological replicates shown as dots. *p<0.05, **p<0.01, ***p<0.001 See also Supplemental Figure 5.

To investigate if the synergism observed with synthetic constructs was conserved in endogenous regulatory regions, we tested the activity of previously identified putative *Foxd1* regulatory regions (PP1, PP2, and +37086) that contained **d**GBMs and E-boxes. Similar to our synthetic reporters, Gli3 alone induced luciferase expression in *Foxd1* PP1, PP2, and +37086 (Fig. 6B-D). Hand2 alone induced activity of PP1 and +37086 but did not significantly increase luciferase activity of PP2 relative to the control. Similar to the output observed with synthetic constructs, co-expression of Gli3 and Hand2 elicited significant and synergistic outputs at all three putative regulatory elements containing endogenous **d**GBMs (Fig. 6B-D; light green hatched bars). This surprising synergism between Gli3 and Hand2 was further confirmed *in vitro* by examining *Foxd1* expression in O9-1 cells, where the presence of Gli3 and Hand2 culminated in synergistic expression of *Foxd1* (Supp. Fig. S5B).

To confirm that this synergism was dependent upon the presence of both a **d**GBM and E-box, we performed site-directed mutagenesis. Mutation of either the **d**GBM or E-box sequence eliminated synergistic output in *Foxd1* endogenous putative regulatory regions (Fig. 7A; Supp. Fig. S6). Furthermore, to determine if the central ‘T’ which we used to define **d**GBMs was causative for the synergistic output, we mutated the ‘T’ in the PP2 putative regulatory region to an ‘A’, resembling a **c**GBM. This single base-pair ‘T>A’ change significantly increased affinity of Gli3 for the GBM and abolished the synergistic luciferase output when Gli3 and Hand2 were co-expressed (Fig. 7B-E). Together, these data support a novel, tissue-specific transcriptional mechanism in which Gli3 and Hand2 utilize low-affinity **d**GBM and E-boxes to promote synergistic activation of *Foxd1* (and likely other MNP targets) outside of a Hh gradient (Fig. 7F, G).

**Figure 7.**
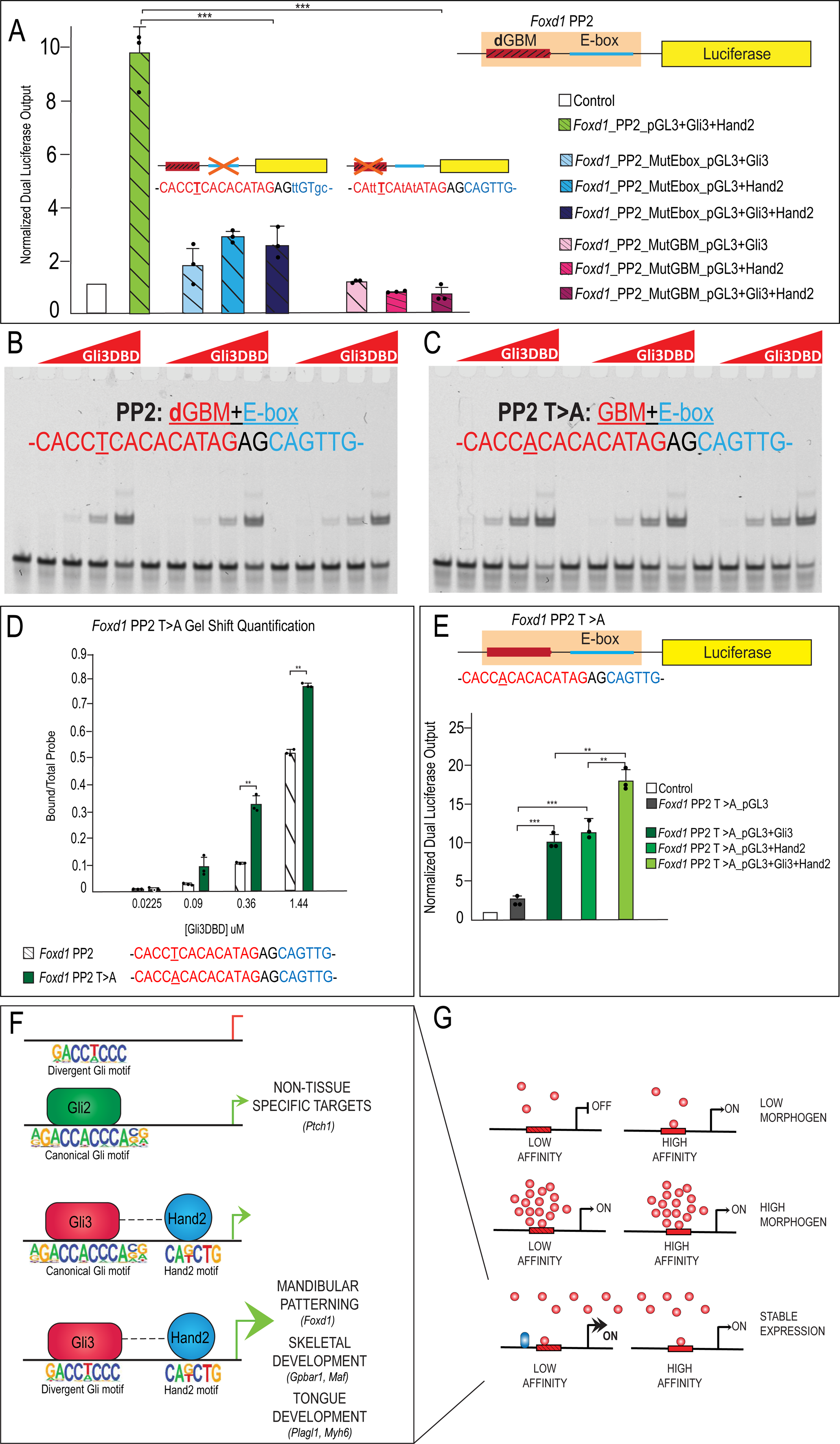
A low affinity **d**GBM occupancy and E-box are required for transcriptional synergism (A) Luciferase reporter activity of mutant GBM or E-box motifs from *Foxd1* PP2 showing mutation of E-box or GBM abolishes synergistic activation. (B-C) EMSA for Gli3DBD binding affinity of (C) endogenous *Foxd1* PP2 or (D) T>A mutant *Foxd1* PP2. (D) Quantification of (B) and (C) showing increased Gli3DBD binding affinity of endogenous *Foxd1* PP2 (white hatched) compared to T>A mutant *Foxd1* PP2 (green). (E) Increased luciferase reporter activity when T>A change is made within *Foxd1* PP2. (F-G) Model of Gli3-Hand2 specific cooperation at low affinity dGBMs drives skeletal and glossal GRN in regions of the MNP outside of the highest Shh ligand concentration. Data are expressed as mean + SD. Luciferase data have biologic replicates shown as dots. *p<0.05, **p<0.01, ***p<0.001. See also Supplemental Figure 6.

## DISCUSSION

Substantial evidence has long supported the idea that the Hh signaling pathway utilizes a morphogen gradient to convey a threshold of activation responses necessary to pattern tissues throughout the embryo (Dessaud et al. 2008). While the concept of a morphogen gradient has been supported by several biochemical and genetic studies, a significant gap remains in understanding the mechanisms of how cells perceive and transduce morphogens. This knowledge gap is especially evident within the developing craniofacial complex, where despite requiring a localized, epithelial Hh source, neither a *Gli* gradient nor a primary requirement of a single Gli (e.g., Gli2 or Gli3) is apparent within facial prominences. In this study, we have uncovered a unique mechanism used in the developing mandible that produces synergistic target gene responses outside of a traditional morphogen gradient by utilizing regulatory elements containing low-affinity GBMs that integrate input from a tissue-specific binding partner. Specifically, our results establish a novel relationship between Gli3 and Hand2, in which these factors synergize at low affinity ‘divergent’ GBMs (**d**GBMs) for a subset of target genes important for key processes in mandibular development including patterning, skeletogenesis and glossogenesis. To our knowledge, this is the first study to identify and explore variable levels of Gli-dependent transcriptional activity across a field of cells as a mechanism for generating cellular identities in the developing face.

### Low-affinity GBMs function as important transcriptional determinants

TF binding site affinity is one mechanism utilized by cells in other tissues to produce graded threshold responses (Driever et al. 1989; Oosterveen et al. 2012; Peterson et al. 2012). The established model states that target genes within a high concentration of the morphogen gradient are activated through low-affinity sites (Jiang and Levine 1993), whereas those exposed to lower morphogen concentrations utilize high-affinity sites (Ip et al. 1992). Despite the validation of this idea in many contexts, regulation of several Hh targets are inconsistent with this model. For example, in the *Drosophila* imaginal disc, *ptc* is restricted to the highest Hh threshold and is regulated by high-affinity canonical GBMs, whereas, *dpp* is expressed broadly throughout the Hh gradient and is regulated by low-affinity non-canonical GBMs (Wang and Holmgren 1999; Parker et al. 2011). Previous ChIP studies in the developing limb have reported that while 55% of Gli binding regions contained a high-affinity GBM, the remaining 45% of regions contained a low-affinity GBM or no GBM.

Interestingly, low-affinity GBMs are strongly conserved across both tissues and species (Vokes et al. 2008; Parker et al. 2011). Furthermore, the same study also reported that a small number of Gli binding regions contained limb-specific variants of the GBM, supporting previous reports that low-affinity motifs are absolutely critical to confer spatially distinct gene expression (Jiang and Levine 1993; Lebrecht et al. 2005; Vokes et al. 2008). Our findings are the first to report how GBM affinity is utilized in a craniofacial context. In the face, we identified low-affinity, divergent binding sites that were necessary and sufficient to drive robust gene expression required for mandibular development (Fig. 7F, G). Interestingly, as no discernable concentration gradient of *Gli2/Gli3* in the developing mandible exists, the utilization of these low affinity divergent sites is likely not dictated by a graded Hh signal. Therefore, our study provides a mechanism for activating transcription networks necessary for mandibular patterning, skeletogenesis and glossogenesis independent of a Shh morphogen gradient.

Previous studies examining Gli binding in the limb and central nervous system (CNS) have identified E-boxes within Gli ChIP-seq peaks. *De novo* motif analysis revealed an E-box enriched in limb Gli binding regions with or without a high-affinity GBM (Vokes et al. 2008). At the time, the significance of the E-box to Gli3 transcriptional activity was unknown. In the developing CNS, an E-box was the second ranked motif identified in Gli1 ChIP-seq peaks (Lee et al. 2010). Mutational analyses determined these E-boxes had varying (context-specific) effects on Gli-mediated transcription, sometimes conferring no affect, while in other cases reducing Gli1-responsiveness (Lee et al. 2010). Our studies significantly advance these findings by demonstrating that co-utilization of GBMs and E-boxes allows Gli TFs to utilize lower affinity sites and produce synergistic transcriptional outputs. Furthermore, our mutational analyses revealed that a single base-pair substitution (“A” with a “T” at the central 5^th^ residue) was sufficient to convey both affinity and synergism. Interestingly, similar divergent, low-affinity GBMs with a medial “T” were previously reported in *Drosophila* within the *dpp* enhancer (Parker et al. 2011). In light of the *dpp* expression pattern, which is broad and found throughout the Hh gradient, it is tempting to speculate that the presence of this medial “T” and subsequent low-affinity GBM could be an evolutionarily conserved mechanism used to generate variable levels of Hh target gene expression independent of a Hh threshold and distinct activator and repressor Gli isoforms.

### Interactions with other TFs control context specific functions of Gli TFs in the face

While traditional descriptions of Gli-mediated Hh signal transduction do not include the requirement of binding partners, there is an established precedence for this concept. A number of TFs have been implicated as partners capable of interacting with Gli TFs and subsequently modulating Gli transcriptional activity. For example, Gli and Zic proteins were previously reported to physically interact through their zinc-finger domains to regulate subcellular localization and transcriptional activity important during neural and skeletal development (Brewster et al. 1998; Koyabu et al. 2001; Zhu et al. 2008). The pluripotency factor Nanog was also reported to physically interact with enhancer-bound Gli proteins to reduce the transcriptional response of cells to a Hh stimulus (Li et al. 2016). The Sox family of TFs have also been implicated in associating with Gli proteins to modulate transcriptional responses in various tissues (Peterson et al. 2012; Tan et al. 2018). Within the developing neural tube, Sox2 was determined to have a significant number of overlapping target genes, as Gli1 and Gli1/Sox2-bound CRMs were shown to induce Shh target gene expression (Peterson et al. 2012). Furthermore, Sox9 and Gli directly and cooperatively regulate several genes important in chondrocyte proliferation (Tan et al. 2018). Finally, recent studies have revealed that the bHLH TF Atoh1 synergizes with Gli2 to activate a medulloblastoma transcriptional network (Yin et al. 2019).

While several previous studies have reported interactions between Hand2 and the Gli TFs in the limb and in establishing left-right asymmetry, the mechanistic relationship appears to be tissue-specific and facets of this relationship still remain elusive. For example, Hand2 is believed to function downstream of Shh during establishment of left-right asymmetry (Olson and Srivastava 1996), while Hand2 is believed to regulate *Shh* expression in the developing limb (Charite et al. 2000; Fernandez-Teran et al. 2000; Yelon et al. 2000; McFadden et al. 2002). Interestingly, in the context of Hand2 acting upstream of Shh, previous studies suggested this to be a DNA-binding-independent effect, (McFadden et al. 2002) and propose that protein-protein interactions or dimer equilibrium can target Hand TFs to regulatory regions (Firulli 2003).

Our work identified a novel relationship between Gli3 and Hand2 that is both unique to the tissue of origin (mandible) and the nature of the interaction (physical interaction, DNA-dependence). First, our RNA-seq analyses on *Gli2^f/f^;Gli3^f/f^;Wnt1-Cre* and *Hand2^f/f^-Wnt1-Cre* mandibular tissue did not reveal any changes in *Hand2* or *Shh* expression, respectively, suggesting that unlike the relationship in the limb or in establishing polarity, there was not a cross-regulatory relationship between Shh and Hand2 in the mandible. Second, our site directed mutagenesis experiments suggested that DNA-binding at some level is required for the Gli3/Hand2 synergism in the MNP, as opposed to the posited DNA-independent mechanism in the limb. Interestingly, the orientation and spacing of E-box and GBMs was not conserved, suggesting flexibility in the architecture underlying Gli3 and Hand2 co-regulatory interactions. The presence of additional TF motifs found in close proximity to GBMs, together with the established knowledge that Gli can interact with a number of other TFs, suggests that a larger protein complex may be at work. Furthermore, the cadre of proteins in this complex could vary depending upon the particular genomic locus and the role it plays regulating transcription either positively or negatively. Future studies will address the role, if any, these other proteins play in modulating Gli transcriptional output in the developing craniofacial complex.

### Gli3 functions as an activator within the developing craniofacial complex

In general, there are two accepted mechanisms for positive Gli-mediated transcriptional regulation: activation and de-repression (Falkenstein and Vokes 2014). Activation refers to the full-length GliA isoform binding regulatory regions of target genes and driving gene expression. Gli1 and Gli2 play the predominant role in activating transcription (Ding et al. 1998; Matise et al. 1998; Park et al. 2000; Stamataki et al. 2005), and are believed to function within the highest concentrations of the Hh gradient (Pan et al. 2006). In the human face, loss of Gli2 has been associated with several craniofacial anomalies presenting with loss-of-function Hh phenotypes such as microcephaly, hypotelorism and a single central incisor (Roessler et al. 2003). Interestingly, mutations in Gli2 have no reported effects on the mandible.

De-repression is the second major mechanism of Hh signal transduction. In this case, targets silenced by the GliR require alleviation of this repression for expression. Subsequent activation can then occur from either GliA or additional TFs. GliR function is primarily carried out by Gli3 and is indispensable for proper Hh-dependent patterning (Litingtung and Chiang 2000; Wang et al. 2000; Oosterveen et al. 2012; Lex et al. 2020). Recent studies in the limb, which is dependent upon Gli3R for proper patterning, have shown that not all GBMs are created equal. While some GBMs appear to be completely dependent on a Hh input, others remain ‘stable’ with Gli3 occupancy occurring independent of the morphogen (Lex et al. 2020). However, to date, no classification system of GBM utilization in the face has been established. In the face, Gli3R is also the predominant repressor for facial patterning, as loss of Gli3 has been associated with several human craniofacial anomalies presenting with gain-of-function Hh phenotypes including Greig cephalopolysyndactyly (Vortkamp et al. 1991; Vortkamp et al. 1992; Hui and Joyner 1993; Wild et al. 1997). Our current study reveals a previously unappreciated role for Gli3A in craniofacial development. Our genetic, biochemical and genomic data suggest Gli3/Hand2 complexes are specifically required to initiate patterning of the MNP and skeletogenic/glossogenic transcriptional networks. Several possibilities exist to explain why Hand2-dependent synergistic activation of targets may be unique to Gli3. First, while Gli3R is highly stable, Gli3A is reportedly not as stable as Gli2A (Pan et al. 2006; Humke et al. 2010; Wen et al. 2010). Association with Hand2 (and possibly other TFs in complex) may stabilize Gli3A, preventing degradation and allowing the isoform to function more efficiently. A second possibility is that Gli2A may predominantly utilize high-affinity GBMs to activate pathway targets, while Gli3A (when in complex with Hand2) predominantly utilizes low-affinity GBMs to activate tissue-specific targets independent of Hh concentration. Determining if these possibilities exist within the mandible, and within other craniofacial prominences, is one aspect of our ongoing work.

In closing, our results reveal a novel transcriptional mechanism for Gli signal transduction in the developing craniofacial complex outside of the traditional graded Hh signaling domains. Our data, compared to that in other organ systems, highlight the diversity of mechanisms utilized by Gli TFs across different tissues. As an organ system, the craniofacial complex is unique because it originates from facial prominences that constitute distinct developmental fields, in both cell content and transcriptional profiles. Thus, as Hand2 is only expressed in the mandibular prominence, our data pose the interesting possibility that facial prominences use unique, prominence-specific Gli partners to transduce Gli signals during craniofacial development. Furthermore, our data suggests sequence variation with in GBMs, may also contribute to tissue-specific Gli transcriptional output. The discovery that a single base-pair within GBMs can relay significant transcriptional activity may lend new insight into examining genetic mutations in human patients with craniofacial anomalies.

## MATERIALS AND METHODS

### Mouse strains

The *Wnt1-Cre*, *Hand2^fl^* (Stock No 027727), and *Gli3^fl^* (Stock No 008873) mouse strains were purchased from Jackson Laboratory. *Gli2^f/f^* mice were provided by Dr. Alexandra Joyner at Memorial Sloan-Kettering Cancer Center. As described in PMID 18501887, conditional deletion of *Hand2* using Wnt1-Cre is embryonic lethal ∼E12 due to loss of norepinephrine. To rescue this phenotype and for investigation of *Hand2^f/f^;Wnt1-Cre* mutants at later embryonic stages, beginning at embryonic day 8 (E8), pregnant dams were fed water containing 100 μg/ml L-phenylephrine, 100 μg/ml isoproterenol, and 2 mg/ml ascorbic acid. All mice were maintained on a CD1 background. Both male and female mice were used. A maximum of 4 adult mice were housed per cage, and breeding cages housed 1 male paired with up to 2 females. All mouse usage was approved by the Institutional Animal Care and Use Committee (IACUC) and maintained by the Veterinary Services at Cincinnati Children’s Hospital Medical Center. N≥5 biologic replicates (biologically distinct samples) for each genotype shown.

### Embryo collection and tissue preparation

Timed mouse matings were performed, with noon of the day a vaginal plug was discovered designated as embryonic day (E) 0.5. Embryos were harvested between E10.5-18.5, collected in PBS, and fixed in 4% paraformaldehyde (PFA) overnight at 4°C, unless otherwise noted. For paraffin embedding, embryos were dehydrated through an ethanol series, washed in xylene, and embedded in paraffin.

### Skeletal preparations

For skeletal preparations, E18.5 embryos were immersed in hot water before skin and soft tissue were removed. Embryos were then stored in acetone for 48 hours, 0.015% alcian blue for 24 hours, washed with ethanol for 24 hours, immersed in 1% fresh KOH for 24-31 hours, then stained with 0.005% alizarin red for 15 hours, and transitioned through a series of glycerol dilutions.

### RNAscope in situ hybridization

Paraffin-embedded embryos were cut at 5µm, and staining was performed with the RNAscope Multiplex Fluorescent Kit v2.0 according to the manufacturer’s instructions. Briefly, sections were deparaffinized in xylene, rehydrated through an ethanol series, and antigen retrieval was performed. The following day, probes were hybridized to sections, paired with a fluorophore, and mounted with Prolong Gold after counterstaining with DAPI. *Shh, Gli2, Gli3, Hand2, Foxd1, Myh6, Gpbar1, Maf,* and *Plagl1* probes for the assay were designed and synthesized by Advanced Cell Diagnostics. RNAScope experiments were performed on at least N≥3 biological replicates for each probe.

### RNA Extraction and Reverse Transcription

RNA was extracted from cells using Trizol™-Micro Total RNA Isolation Kit (Invitrogen, 15596026). cDNA was synthesized from up to 2µg of RNA with the High Capacity RNA-to-cDNA Kit (Invitrogen, 4387406).

### Quantitative Real-Time PCR

qRT-PCR was performed in technical (multiple replicates of the same biological sample) triplicate using PowerUP SYBR Green Master Mix (ThermoFisher Scientific, A25742) on Applied Biosystems QuantStudio 3 Real-Time PCR System (ThermoFisher Scientific) for N=3 biological replicates. All genes were normalized to *Gapdh* expression.

### Co-immunoprecipitation

Mandibular prominences were harvested from E10.5 CD-1 embryos, pooled, and lysed in RIPA buffer containing Halt protease inhibitor cocktail. Protein lysate was incubated with Hand2 (polyclonal goat IgG) or control goat IgG primary antibody overnight at 4C with nutation. Dynabeads Protein G were added the next day and incubated with antibody-lysate mixture for 4 hours at 4C on a nutator. Dynabeads Protein G-antibody-antigen complex was washed 3 times using RIPA buffer, and antigens were eluted from the beads in SDS sample buffer by boiling for 5 minutes. N=4 biological replicates of pooled litters.

### Western blotting

For co-immunoprecipitation, eluted products and 10% of the input were separated by SDS-PAGE and transferred to a PVDF membrane for blotting at 4C with Gli3 (polyclonal goat IgG 1:1000, R&D Systems) and Hand2 (polyclonal goat IgG or mouse monoclonal IgG1 1:1000) primary antibodies. Detection of primary antibodies was performed using infrared-conjugated secondary antibodies (donkey anti-goat or goat anti-mouse IRDye 800CW, LICOR) and acquired using a LICOR infrared scanner. For plasmid verification, Flag primary antibody (monoclonal M2 mouse IgG1) and enhanced chemiluminescence assay (Amersham ECL Primer, GE Healthcare Life Science) were used for detection.

### Single cell RNA-sequencing

Mandibles from E11.5 or E13.5 wildtype CD1 mouse embryos were quickly dissected in ice-cold PBS and minced to a fine paste. Cells were dissociated into a single-cell suspension and sequenced using NovaSeq 6000 and the S2 flow cell. 12.5mg of tissue was placed in a sterile 1.5mL tube containing 0.5mL protease solution containing 125 U/ml DNase and *Bacillus Licheniformis* (3mg/ml for e11.5 sample and 5mg/ml for e13.5 sample). The samples were incubated at 4C for a total of 10 minutes, with trituration using a wide boar pipette tip every minute after the first two. Protease was inactivated using ice-cold PBS containing 0.02% BSA and filtered using 30µM filter. The cells were pelleted by centrifugation at 200G for 4 minutes and resuspended in 0.02% BSA in PBS. Cell number and viability were assessed using a hemocytometer and trypan blue staining. 9,600 cells were loaded onto a well on a 10x Chromium Single Cell instrument (10X Genomics) to target sequencing of 6,000 cells. Barcoding, cDNA amplification, and library construction were performed using the Chromium Single Cell 3’ Library & Gel Bead Kit v3. Post cDNA amplification and cleanup was performed using SPRI select reagent (Beckman Coulter, Cat# B23318). Post cDNA amplification and post library construction quality control was performed using the Agilent Bioanalyzer High Sensitivity kit (Agilent 5067-4626). Libraries were sequenced using a NovaSeq 6000 and the S2 flow cell. Sequencing parameters used were: Read 1, 28 cycles; Index i7, 8 cycles; Read 2, 91 cycles, producing about 300 million reads. The sequencing output data was processing using CellRanger (http://10xgenomics.com) to obtain a gene-cell data matrix.

### Chromatin Immunoprecipitation

Individual ChIP-seq experiments were carried out on pooled embryonic tissue collected in ice-cold PBS. Dissected tissues were immediately fixed in 1% formaldehyde/PBS for 20 minutes at room temp followed by glycine quench (125 mM). ChIP procedures were performed as previously described (Peterson et al. 2012 and Osterwalder et al. 2014). All ChIP experiments were performed using mouse monoclonal anti-FLAG M2 antibody (Sigma-Aldrich). A mock control ChIP sample was made by performing ChIP on tissues isolated from wild-type embryos.

### RNA-sequencing

MNPs were dissected from E10.5 embryos, using at three biologic samples. RNA was prepared for RNA-seq using Invitrogen RNAqueous™-Micro RNA Isolation Kit (AM1931). Sequencing was carried out in 150 bp paired-end reads using the Illumina HiSeq2500 system.

### ATAC-seq

Individual E11.5 MNP’s were collected from wild-type embryos and immediately snap frozen in liquid nitrogen. Nuclei were isolated by incubating in homogenization buffer (250 mM sucrose; 25 mM KCl; 5 mM MgCl2; 20 mM Tricine-KOH; 1mM EDTA; and 1% IGEPAL) for 30 minutes at 4C with shaking (800 rpm). Cell nuclei were counterstained with Trypan Blue and counted. Approximately 5X10^4^ nuclei were processed for ATAC-seq as previously described (Buenrostro et al. 2015). DNA libraries were sequenced on NextSeq550 (Illumina) to generate 75 bp paired-end reads.

### Protein purification and EMSA

Coding regions for all protein fragments used for EMSA were cloned in-frame with an N-terminal 6xHis-tag in the pET14b vector (Novagen) and expressed in BL21 cells. The mouse E47 (E47L) isoform of the Tcf3 protein containing the bHLH domain (amino acids 271 to 648), the mouse Gli3 (Gli3DBD) protein containing the 5 zinc fingers in its DNA binding domain (amino acids 465-648), and the full-length mouse Hand2 (Hand2FL) protein (amino acids 1-217) were purified under denaturing conditions via Ni-chromatography and refolded in Native lysis buffer while on Ni-beads as described previously (Witt et al. 2010; Zhang et al. 2019). Expression of each protein was confirmed via coomassie staining, and protein concentrations were measured via Bradford Assay. Probes were generated as previously described by annealing a 5’IREdye-700 labeled oligo from IDT with the following sequence 5’-CTATCGTAGACTTCG-3’ to each oligo listed below and filling in via a Klenow reaction (Uhl et al. 2016). EMSAs were performed as previously described with the following modification to allow homodimer and heterodimer exchange between bHLH proteins (E47 and Hand2): binding reactions were incubated at 37°C for 40 minutes before allowing each reaction to cool to room temperature and incubating with DNA probes for an additional 15 minutes prior to separation on a native SDS gel (Uhl et al. 2010; Uhl et al. 2016). All EMSAs were imaged using a LICOR Clx scanner.

### Plasmid Constructs

Luciferase reporter constructs were generated by cloning putative enhancer fragments into the pGL3-promoter luciferase reporter plasmid. Hand2 and Gli3, were all cloned into a p3XFlag CMV 7.1 plasmid.

### Luciferase Reporter Assay

O9-1 neural crest cells were maintained in conditioned media collected from mitomycinC-inactivated fibroblasts supplemented with LIF and b-FGF. Cells were co-transfected in triplicate with the appropriate luciferase reporter plasmid, a Renilla control plasmid, and a combination of plasmids expressing Gli2, Gli3, or Hand2 using Lipofectamine 3000. Cells were harvested 24 hours after transfection, and luciferase activity was determined using the Dual Luciferase Reporter Assay System (Promega) and the GLOMAX luminometer. N≥3 biological replicates performed in technical triplicate for each condition.

### Bulk RNA-seq Analysis

Paired-end reads were mapped to mm10 genome and transcript abundance was determined using Strand NGS. Differential expression was determined using DESeq2 within Strand NGS.

### Single Cell RNA-Seq Analysis

Raw reads were sequenced using 10x v2 chemistry for two samples E11.5 and E13.5 MNP. Reads were mapped to mouse transcriptome (mm10) version of the UCSC using Cellranger (https://github.com/10XGenomics/cellranger). Approximately 70% of the reads were confidently mapped to the transcriptome and ∼2500 genes were expressed per sample. Quality control (QC) was carried out where cells with less than ∼1k UMI’s were removed from the quantification analysis. Finally, raw reads were quantified into a raw-counts matrix for cells that passed QC.

Raw counts matrix was analyzed using Seurat (v2.3.4) (Stuart et al. 2019). Briefly, all genes expressed in ≥3 cells and cells with at least 200 genes expressed were used for downstream analysis. Quality filtering of cells was done based on number of genes expressed and percent of mitochondrial expression. Followed by filtering, normalization of data was carried out using log2 transform and a global scaling factor. Highly variable genes (HVGs) which exhibit cell-to-cell variation, were selected by marking the outliers on average Expression vs dispersion plot and cell cycle effect was regressed by removing the difference between the G2M and S phase. Next, HVGs were used to perform a linear dimension reduction using principal component analysis (PCA) and top 20 principal components (PCs) were used to cluster cells into respective clusters using graph-based knn clustering approach. Markers for each cluster were obtained using Wilcoxon rank sum test in ‘FindAllMarkers’ function. Cell clusters were annotated to respective cell-types using a-priori knowledge of defined cell-type markers. Finally, clusters were visualized using t-distributed stochastic neighbor embedding (tSNE) a non-linear dimension reduction.

Further, to understand the similarities and differences among cell-types annotated in each sample (E11.5, E13.5 MNP), an integration analysis was performed using Seurat (v3.0) (https://github.com/Brugmann-Lab/Single-Cell-RNA-Seq-Analysis). Quality filtering, normalization, cell-cycle regression was performed as explained above. Feature selection (selecting HVGs) was done using variance stabilizing transform (vst) method as described in Seurat tutorial. Next, dimensionality reduction for both samples together was performed using diagonalized canonical correlation analysis (CCA) followed by L2-normalization and finally searching for mutual nearest neighbors (MNNs). Resulting cell-pairs from MNN were annotated as anchors (‘FindIntegrationAnchors’ function Seurat). Those integration anchors were then used to integrate the samples using ‘IntegrateData’ function in Seurat. After integrating the datasets, PCA was performed on integrated data, top 20 PCs were used for cell clustering using graph-based KNN algorithm and the clusters were visualized uniform manifold approximation projection (UMAP). All the visualization of the single cell data was performed using data visualization functions embedded in Seurat.

### ChIP-seq Analysis

ChIP-seq libraries were prepared according to manufacturers’ instructions and 1X75bp reads were generated on a NextSeq instrument (Illumina). The resulting reads were mapped to mouse genome assembly mm10 (GRCm38/mm10) using bwa (Li and Durbin 2009). Pooled replicates were used to identify potential regulatory regions (Supp. File 3). A final set of peak calls for each factor to use for motif enrichment was determined using bedtools (Quinlan and Hall 2010) to merge biological replicates and identify peaks shared between replicates (Supp. File 4). ChIP-seq peak overlap significance was calculated using the RELI software package (https://github.com/WeirauchLab/RELI) (Harley et al. 2018). TF binding site motif enrichment analyses were performed using the HOMER software package (Heinz et al. 2010) modified to use a log 2-based scoring system and contain mouse motifs obtained from the Cis-BP database, build 1.94d (Weirauch et al. 2014). DNA 8mer counts were calculated by examining the number of times each of the possible 32,896 8mers occurs in the sequences contained within the given ChIP-seq peakset (on either strand, avoiding double-counting for palindromic sequences). Enrichment for particular orientations and spacings between Gli and Hand motifs was performed using the COSMO software package (Narasimhan et al. 2015).

### Statistical Analysis

qPCR and luciferase data are represented as mean+SD. Relative luciferase output was calculated by normalizing raw Luciferase output to Renilla output and comparing this dual luciferase output to a control condition. Statistical significance was determined using Student’s *t* test. *P-*value <0.05 was considered statistically significant. * *p*<0.05, ** *p*<0.01, and *** *p*<0.001.

## ACKNOWLEDGEMENTS

The authors would like to acknowledge the CCHMC DNA Sequencing Core, Gene Expression Core, and Veterinary Services Cores, the Genome Technologies services at The Jackson Laboratory, and the Center for Epigenomics at the University of California San Diego. AZ and RZ thank Jens Stolte for expert technical assistance. We acknowledge support from the National Institute of Health, National Institute for Dental and Craniofacial Research to SAB (R35 DE027557) and KHE (F31 DE027872) and National Institute of General Medical Sciences to KAP (R01 GM124251).

## AUTHOR CONTRIBUTIONS

KHE performed all murine, cell culture, luciferase and bulk RNA-sequencing experiments, participated in bioinformatics analysis, scRNA-seq analysis, figure generation and writing of manuscript. XC and MTW performed ChIP-seq bioinformatics analysis. JSalomone and BG purified protein for and performed EMSAs. PC provided genomic coordinates for CisBP-identified **c**GBMs and performed scRNA-seq bioinformatic analysis. PS provided support for murine and luciferase-based experiments. SKB collected scRNA-seq samples. AZ and RZ performed Hand2 ChIP experiments. JDS and KAP performed GLI3 ChIP-seq and bioinformatic analysis. KAP generated ATAC-seq data. KHE, MTW, BG, KAP and SAB discussed the project and edited the manuscript. SAB formulated the project, wrote the manuscript, edited figures, and analyzed data.

---

**Supplemental Figure 1.**
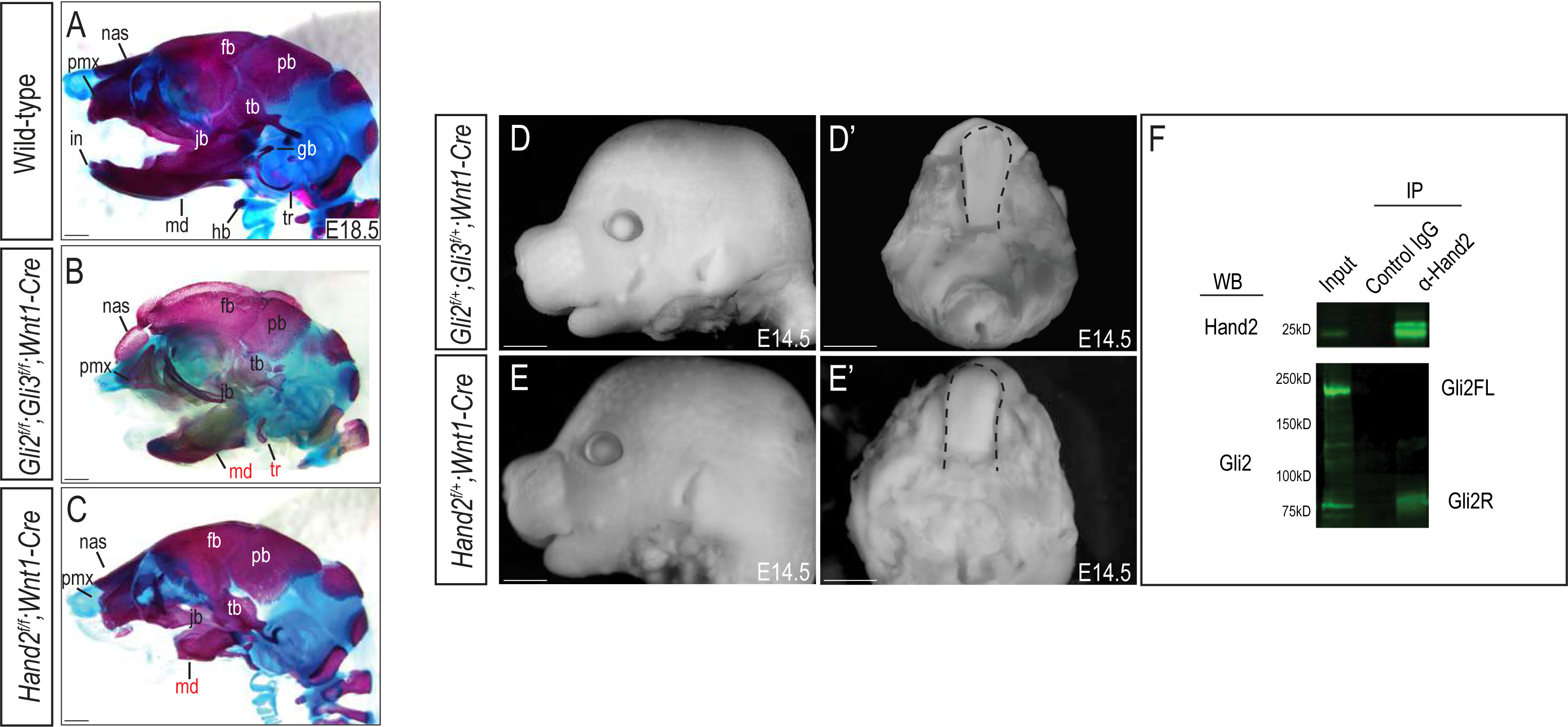
Heterozygotes for Gli or Hand2 in NCCs display no MNP phenotype (A-C) Lateral cranial view of skeletal stains for whole heads of wild-type and *Gli2^f/f^;Gli3^f/f^;Wnt1-Cre* and *Hand2^f/f^;Wnt1-Cre* mutants at E18.5. Abbre-viations: agp, angular process; pmx, pre-maxilla; nas, nasal bone; jb, jugal bone; hb, hyoid bone; fb, frontal bone; pb, parietal bone; tb, temporal bone; g, gonial; tr, tympanic ring; hb, hyoid bone. (D, E) Lateral cranial view and (D’, E’) dorsal mandibular view of *Gli2^f/+^;Gli3^f/+^;Wnt1-Cre* and *Hand2^f/+^;Wnt1-Cre* embryos at E14.5. Dotted black line indicates presence of tongue in both allelic combinations. (F) Co-immunoprecipitation showing interaction between Gli3 and Hand2 within E10.5 MNPs. Scale bar: 1mm.

**Supplemental Figure 2.**
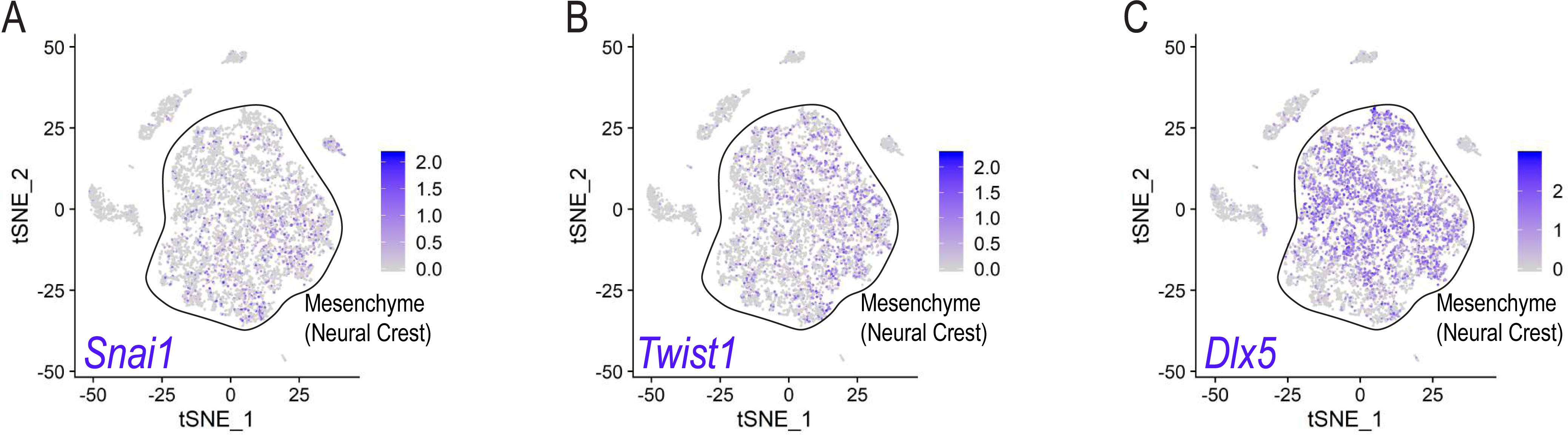
cNCC derivates in the early MNP (A-C) Single cell expression of *Snai1, Twist1*, and *Dlx5* in the E11.5 MNP. Black outline emphasizes the enriched expression of these markers to indicate NCC-derived mesenchyme.

**Supplemental Figure 3.**
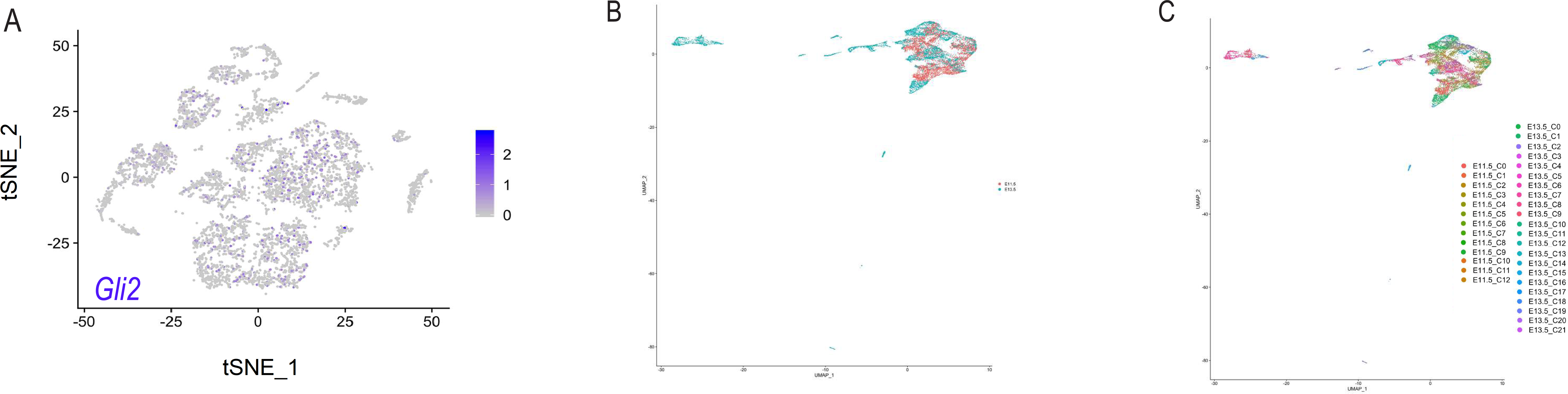

**Supplemental Figure 4.**
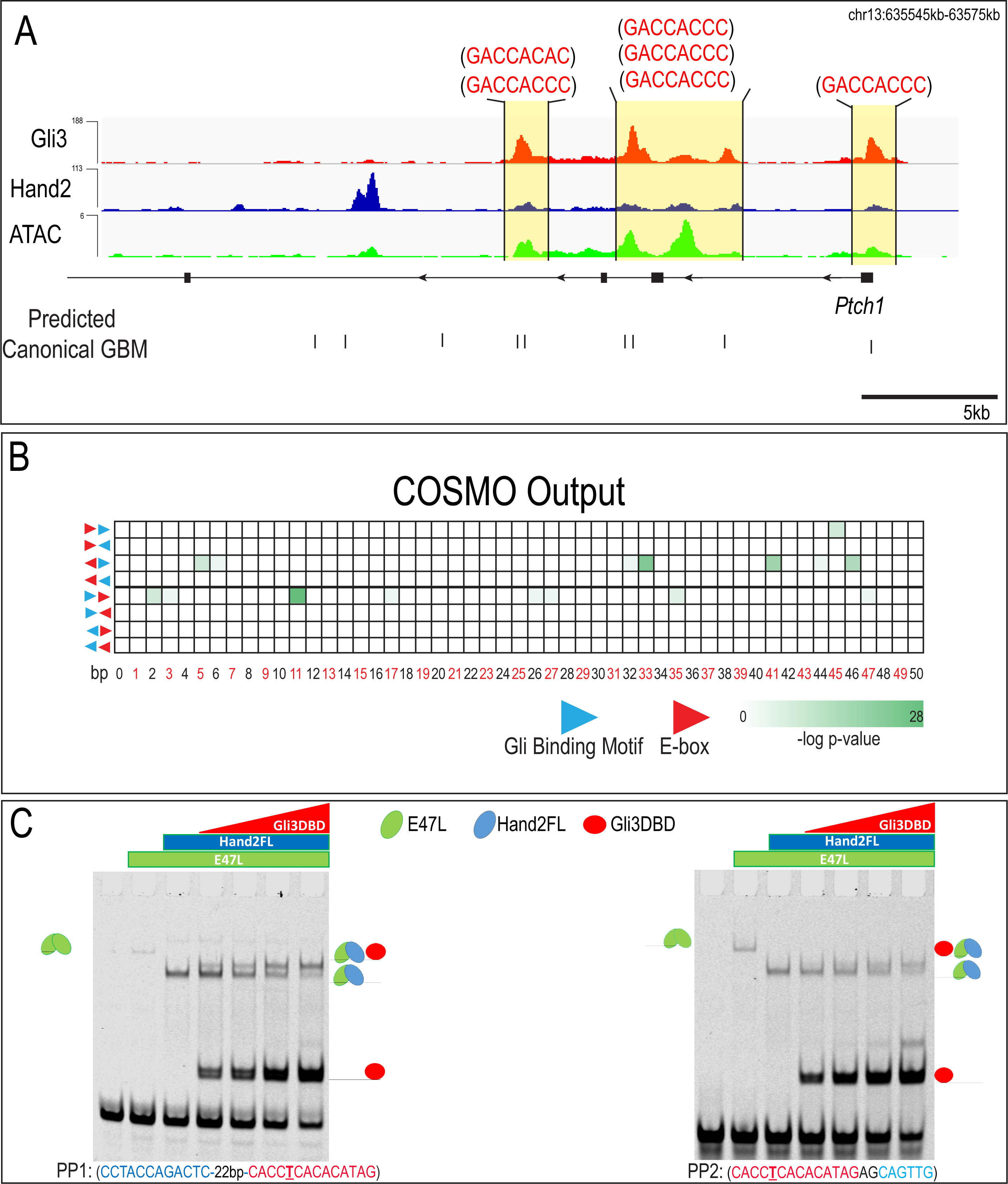
Spacing and orientation constraints were not observed between Gli and Hand2 binding sites. (A) MNP regulatory landscape for *Ptch1*. Predicated **c**GBMs are shown as black tickmarks below the tracks and the associated motif is shown above. (B) Heatmap showing *p*-values associated with COSMO outputs for spacing and orientation of TF binding site enrichment. Number of base pairs separating motifs increase from left to right, orientations and directionality of motifs tested are schematized on the left. No single orientation or spacing was conserved for GBMs and E-boxes within Gli3 and Hand2 overlapping ChIP-seq peaks in the MNP. (C) EMSA for PP1 and PP2 using increasing concentrations of Gli3DBD, Hand2 full-length (FL), and E47L/Tcf3, the Hand2 heterodimerization partner. Increasing concentrations of Gli3DBD results in increased binding of a supershift complex. The observed complexes are illustrated for clarity.

**Supplemental Figure 5.**
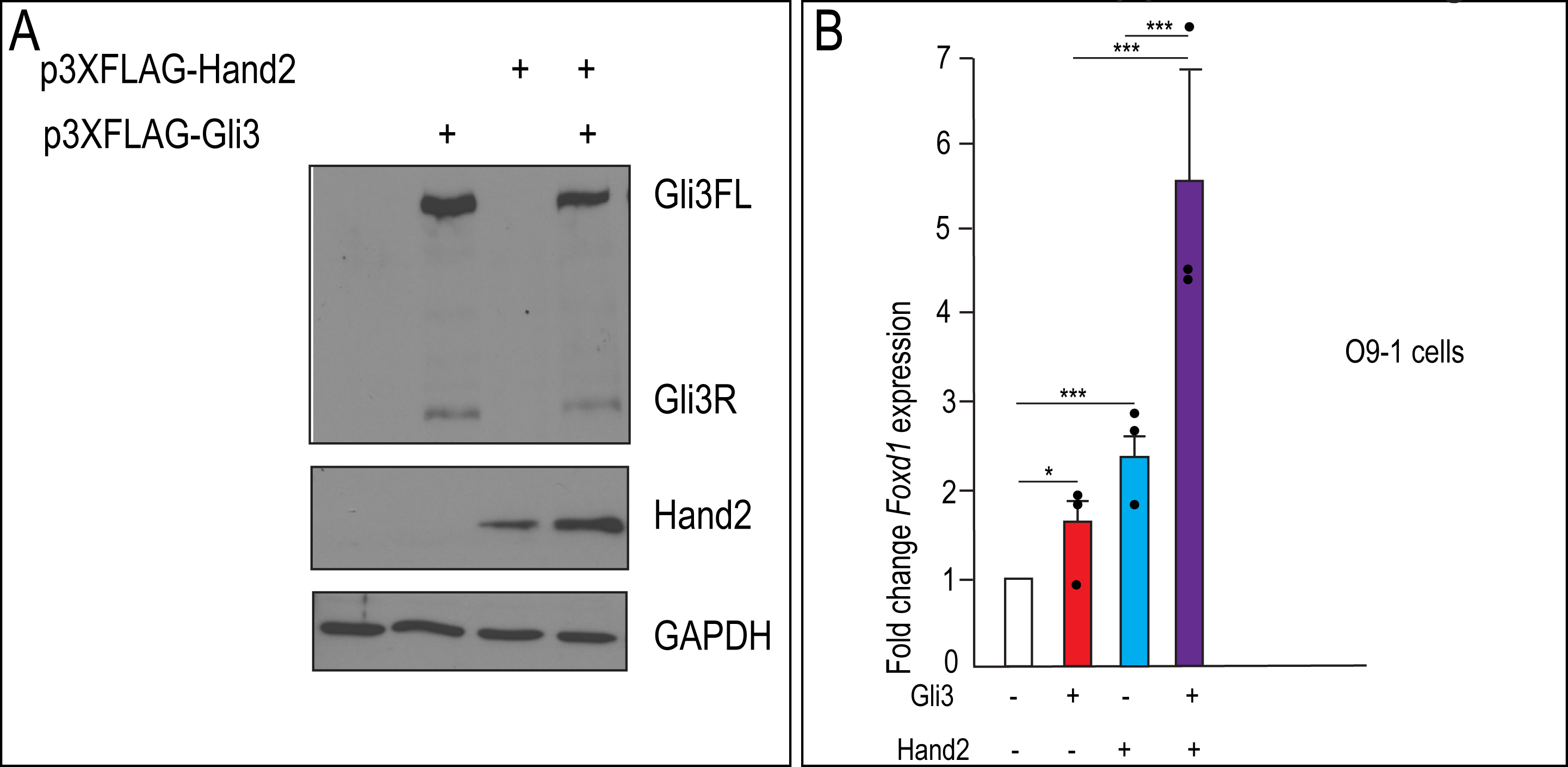
Gli3 and Hand2 co-expression synergistically activates *Foxd1* in vitro (A) Western blot using α-FLAG antibody to detect protein expression from plasmids, p3XFLAG-Hand2 and p3XFlag-Gli3, following transfection into mouse embryonic fibroblasts. (B) RT-qPCR fold change of *Foxd1* after transfection with Gli3, Hand2, or both in O9-1 cells. Data are expressed as mean + SD with biological replicates indicated as dots. *p<0.05, **p<0.01, ***p<0.001.

**Supplemental Figure 6.**
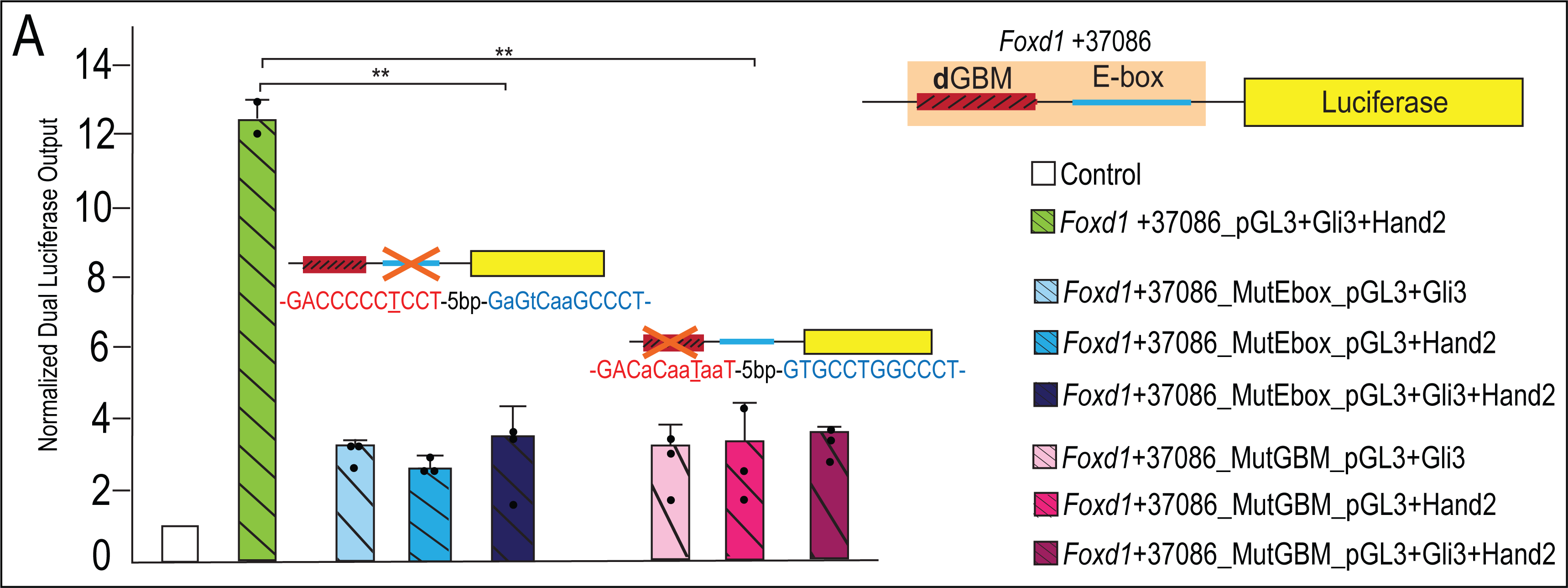
Mutations of **d**GBM or E-box of *Foxd1* +37086 putative enhancer abolish Gli3-Hand2 synergism. (A) Luciferase reporter activity of mutant GBM or E-box motifs from *Foxd1* +37086 showing mutation of E-box or GBM abolishes synergistic activation. Data are expressed as mean + SD with biological replicates indicated as dots. *p<0.05, **p<0.01, ***p<0.001.

**Supplemental Table 1.** Differences in gene expression levels from conditional KO study

| *Gli2<sup>fl</sup>;Gli3<sup>fl</sup>;Wnt1-Cre* |  |  | *Hand2<sup>fl</sup>;Wnt1-Cre* |  |
| --- | --- | --- | --- | --- |
| Gene | Fold Change<sup>a</sup> | Significance<sup>b</sup> | Fold Change<sup>a</sup> | Significance<sup>b</sup> |
| *Hand2* | -1.1 | n.s. | -8.1 | p<0.05 |
| *Gli1* | -3.9 | p<0.05 | -1.0 | n.s. |
| *Gli2* | -1.8 | p<0.05 | -1.1 | n.s. |
| *Gli3* | -2.2 | p<0.05 | -1.1 | n.s. |
| *Ptch1* | -1.8 | p<0.05 | -1.2 | n.s. |
| *Shh* | 1.7 | p<0.05 | 1.0 | n.s. |

<sup>a</sup>Relative to wild-type
<sup>b</sup>not significant; n.s.

**Supplemental Table 2.** smFISH probes provided by ACD Bio for the RNAScope Assay.

| smISH Probes | ACD Bio Product Name | Catalog # | Accession # | Probe region begins | Probe region ends |
| --- | --- | --- | --- | --- | --- |
| Shh | RNAscope® Probe - Mm-Shh | 314361 | NM_009170.3 | 307 | 1197 |
| Ptch1 | RNAscope® Probe - Mm-Ptch1 | 402811 | NM_008957.2 | 2260 | 3220 |
| Gli2 | RNAscope® Probe - Mm-Gli2 | 405771 | NM_001081125.1 | 4409 | 5343 |
| Gli3 | RNAscope® Probe - Mm-Gli3 | 445511 | NM_008130.2 | 716 | 1711 |
| Hand2 | RNAscope® Probe - Mm-Hand2 | 499821 | NM_010402.4 | 2 | 2185 |
| Plagl1 | RNAscope® Probe - Mm-Plagl1 | 462941 | NM_009538.2 | 1117 | 1996 |
| Myh6 | RNAscope® Probe - Mm-Myh6 | 506251 | NM_001164171.1 | 48 | 5162 |
| Gpbar1 | RNAscope® Probe - Mm-Gpbar1 | 318451 | NM_174985.1 | 26 | 955 |
| Maf | RNAscope® Probe - Mm-Maf | 412951 | NM_001025577.2 | 1034 | 2615 |
| Foxd1 | RNAscope® Probe - Mm-Foxd1 | 495501-C3 | NM_008242.2 | 2 | 2220 |

**Supplemental Table 3.**
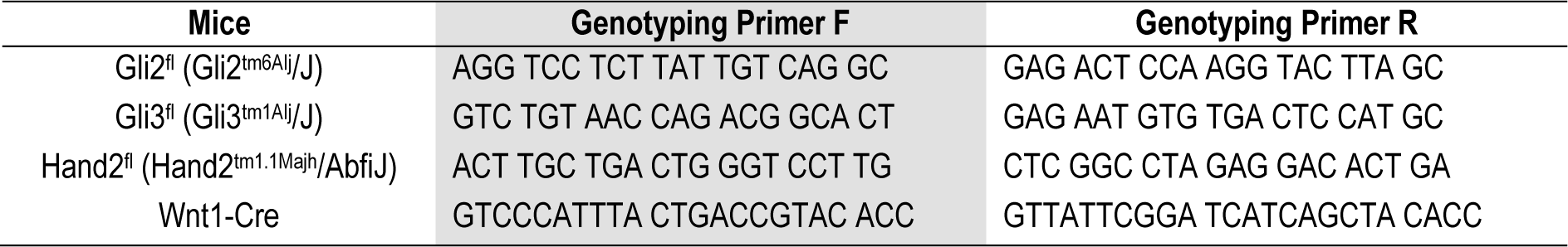
Genotyping primers for transgenic mice.

## REFERENCES

Bai CB, Stephen D, Joyner AL. 2004. All mouse ventral spinal cord patterning by hedgehog is Gli dependent and involves an activator function of Gli3. Dev Cell 6: 103–115.

Balaskas N, Ribeiro A, Panovska J, Dessaud E, Sasai N, Page Karen M, Briscoe J, Ribes V. 2012a. Gene Regulatory Logic for Reading the Sonic Hedgehog Signaling Gradient in the Vertebrate Neural Tube. Cell 148: 273–284.

Balaskas N, Ribeiro A, Panovska J, Dessaud E, Sasai N, Page KM, Briscoe J, Ribes V. 2012b. Gene regulatory logic for reading the Sonic Hedgehog signaling gradient in the vertebrate neural tube. Cell 148: 273–284.

Barron F, Woods C, Kuhn K, Bishop J, Howard MJ, Clouthier DE. 2011. Downregulation of Dlx5 and Dlx6 expression by Hand2 is essential for initiation of tongue morphogenesis. Development (Cambridge, England) 138: 2249–2259.

Berger MF, Bulyk ML. 2009. Universal protein-binding microarrays for the comprehensive characterization of the DNA-binding specificities of transcription factors. Nature protocols 4: 393–411.

Berger MF, Philippakis AA, Qureshi AM, He FS, Estep PW, 3rd, Bulyk ML. 2006. Compact, universal DNA microarrays to comprehensively determine transcription-factor binding site specificities. Nature biotechnology 24: 1429–1435.

Bowers M, Eng L, Lao Z, Turnbull RK, Bao X, Riedel E, Mackem S, Joyner AL. 2012. Limb anterior-posterior polarity integrates activator and repressor functions of GLI2 as well as GLI3. Developmental biology 370: 110–124.

Brewster R, Lee J, Ruizi Altaba A. 1998. Gli/Zic factors pattern the neural plate by defining domains of cell differentiation. Nature 393: 579–583.

Briscoe J, Ericson J. 2001. Specification of neuronal fates in the ventral neural tube. Curr Opin Neurobiol 11: 43–49.

Buenrostro JD, Wu B, Chang HY, Greenleaf WJ. 2015. ATAC-seq: A Method for Assaying Chromatin Accessibility Genome-Wide. Curr Protoc Mol Biol 109: 21 29 21–21 29 29.

Buscher D, Ruther U. 1998. Expression profile of Gli family members and Shh in normal and mutant mouse limb development. Developmental dynamics : an official publication of the American Association of Anatomists 211: 88–96.

Chang CF, Chang YT, Millington G, Brugmann SA. 2016. Craniofacial Ciliopathies Reveal Specific Requirements for GLI Proteins during Development of the Facial Midline. PLoS genetics 12: e1006351.

Chang DT, Lopez A, von Kessler DP, Chiang C, Simandl BK, Zhao R, Seldin MF, Fallon JF, Beachy PA. 1994. Products, genetic linkage and limb patterning activity of a murine hedgehog gene. *Development (Cambridge*, England) 120: 3339–3353.

Charite J, McFadden DG, Olson EN. 2000. The bHLH transcription factor dHAND controls Sonic hedgehog expression and establishment of the zone of polarizing activity during limb development. Development 127: 2461–2470.

Chiang C, Litingtung Y, Lee E, Young KE, Corden JL, Westphal H, Beachy PA. 1996. Cyclopia and defective axial patterning in mice lacking Sonic hedgehog gene function. Nature 383: 407–413.

Cordero D, Marcucio R, Hu D, Gaffield W, Tapadia M, Helms JA. 2004. Temporal perturbations in sonic hedgehog signaling elicit the spectrum of holoprosencephaly phenotypes. The Journal of clinical investigation 114: 485–494.

Dai P, Akimaru H, Tanaka Y, Maekawa T, Nakafuku M, Ishii S. 1999. Sonic Hedgehog-induced activation of the Gli1 promoter is mediated by GLI3. The Journal of biological chemistry 274: 8143–8152.

Dessaud E, McMahon AP, Briscoe J. 2008. Pattern formation in the vertebrate neural tube: a sonic hedgehog morphogen-regulated transcriptional network. *Development (Cambridge*, England) 135: 2489–2503.

Dessaud E, Ribes V, Balaskas N, Yang LL, Pierani A, Kicheva A, Novitch BG, Briscoe J, Sasai N. 2010. Dynamic assignment and maintenance of positional identity in the ventral neural tube by the morphogen sonic hedgehog. PLoS Biol 8: e1000382.

Dessaud E, Yang LL, Hill K, Cox B, Ulloa F, Ribeiro A, Mynett A, Novitch BG, Briscoe J. 2007. Interpretation of the sonic hedgehog morphogen gradient by a temporal adaptation mechanism. Nature 450: 717–720.

Ding Q, Motoyama J, Gasca S, Mo R, Sasaki H, Rossant J, Hui CC. 1998. Diminished Sonic hedgehog signaling and lack of floor plate differentiation in Gli2 mutant mice. *Development (Cambridge*, England) 125: 2533–2543.

Driever W, Thoma G, Nusslein-Volhard C. 1989. Determination of spatial domains of zygotic gene expression in the Drosophila embryo by the affinity of binding sites for the bicoid morphogen. Nature 340: 363–367.

Echelard Y, Epstein DJ, St-Jacques B, Shen L, Mohler J, McMahon JA, McMahon AP. 1993. Sonic hedgehog, a member of a family of putative signaling molecules, is implicated in the regulation of CNS polarity. Cell 75: 1417–1430.

Falkenstein KN, Vokes SA. 2014. Transcriptional regulation of graded Hedgehog signaling. Seminars in cell & developmental biology 33: 73–80.

Fernandez-Teran M, Piedra ME, Kathiriya IS, Srivastava D, Rodriguez-Rey JC, Ros MA. 2000. Role of dHAND in the anterior-posterior polarization of the limb bud: implications for the Sonic hedgehog pathway. Development 127: 2133–2142.

Firulli AB. 2003. A HANDful of questions: the molecular biology of the heart and neural crest derivatives (HAND)-subclass of basic helix-loop-helix transcription factors. Gene 312: 27–40.

Funato N, Kokubo H, Nakamura M, Yanagisawa H, Saga Y. 2016. Specification of jaw identity by the Hand2 transcription factor. Sci Rep 6: 28405.

Hallikas O, Palin K, Sinjushina N, Rautiainen R, Partanen J, Ukkonen E, Taipale J. 2006. Genome-wide prediction of mammalian enhancers based on analysis of transcription-factor binding affinity. Cell 124: 47–59.

Harley JB, Chen X, Pujato M, Miller D, Maddox A, Forney C, Magnusen AF, Lynch A, Chetal K, Yukawa M et al. 2018. Transcription factors operate across disease loci, with EBNA2 implicated in autoimmunity. Nature genetics 50: 699–707.

Hebrok M, Kim SK, St Jacques B, McMahon AP, Melton DA. 2000. Regulation of pancreas development by hedgehog signaling. *Development (Cambridge*, England) 127: 4905–4913.

Heinz S, Benner C, Spann N, Bertolino E, Lin YC, Laslo P, Cheng JX, Murre C, Singh H, Glass CK. 2010. Simple Combinations of Lineage-Determining Transcription Factors Prime cis-Regulatory Elements Required for Macrophage and B Cell Identities. Molecular Cell 38: 576–589.

Helms JA, Kim CH, Hu D, Minkoff R, Thaller C, Eichele G. 1997. Sonic hedgehog participates in craniofacial morphogenesis and is down-regulated by teratogenic doses of retinoic acid. Developmental biology 187: 25–35.

Hong E, Di Cesare PE, Haudenschild DR. 2011. Role of c-Maf in Chondrocyte Differentiation: A Review. Cartilage 2: 27–35.

Hu D, Marcucio RS, Helms JA. 2003. A zone of frontonasal ectoderm regulates patterning and growth in the face. Development 130: 1749–1758.

Hui CC, Joyner AL. 1993. A mouse model of greig cephalopolysyndactyly syndrome: the extra-toesJ mutation contains an intragenic deletion of the Gli3 gene. Nature genetics 3: 241–246.

Hui CC, Slusarski D, Platt KA, Holmgren R, Joyner AL. 1994. Expression of three mouse homologs of the Drosophila segment polarity gene cubitus interruptus, Gli, Gli-2, and Gli-3, in ectoderm- and mesoderm-derived tissues suggests multiple roles during postimplantation development. Developmental biology 162: 402–413.

Humke EW, Dorn KV, Milenkovic L, Scott MP, Rohatgi R. 2010. The output of Hedgehog signaling is controlled by the dynamic association between Suppressor of Fused and the Gli proteins. Genes & development 24: 670–682.

Ip YT, Park RE, Kosman D, Bier E, Levine M. 1992. The dorsal gradient morphogen regulates stripes of rhomboid expression in the presumptive neuroectoderm of the Drosophila embryo. Genes Dev 6: 1728–1739.

Ishii M, Arias AC, Liu L, Chen YB, Bronner ME, Maxson RE. 2012. A stable cranial neural crest cell line from mouse. Stem cells and development 21: 3069–3080.

Jaeger J, Surkova S, Blagov M, Janssens H, Kosman D, Kozlov KN, Manu, Myasnikova E, Vanario-Alonso CE, Samsonova M et al. 2004. Dynamic control of positional information in the early Drosophila embryo. Nature 430: 368–371.

Jeong J, Mao J, Tenzen T, Kottmann AH, McMahon AP. 2004. Hedgehog signaling in the neural crest cells regulates the patterning and growth of facial primordia. Genes & development 18: 937–951.

Jiang J, Levine M. 1993. Binding affinities and cooperative interactions with bHLH activators delimit threshold responses to the dorsal gradient morphogen. Cell 72: 741–752.

Kinzler KW, Vogelstein B. 1990. The GLI gene encodes a nuclear protein which binds specific sequences in the human genome. Molecular and cellular biology 10: 634–642.

Koyabu Y, Nakata K, Mizugishi K, Aruga J, Mikoshiba K. 2001. Physical and functional interactions between Zic and Gli proteins. The Journal of biological chemistry 276: 6889–6892.

Kulakovskiy IV, Medvedeva YA, Schaefer U, Kasianov AS, Vorontsov IE, Bajic VB, Makeev VJ. 2013. HOCOMOCO: a comprehensive collection of human transcription factor binding sites models. Nucleic acids research 41: D195–202.

Lan Y, Jiang R. 2009. Sonic hedgehog signaling regulates reciprocal epithelial-mesenchymal interactions controlling palatal outgrowth. Development 136: 1387–1396.

Lebrecht D, Foehr M, Smith E, Lopes FJ, Vanario-Alonso CE, Reinitz J, Burz DS, Hanes SD. 2005. Bicoid cooperative DNA binding is critical for embryonic patterning in Drosophila. Proc Natl Acad Sci U S A 102: 13176–13181.

Lee EY, Ji H, Ouyang Z, Zhou B, Ma W, Vokes SA, McMahon AP, Wong WH, Scott MP. 2010. Hedgehog pathway-regulated gene networks in cerebellum development and tumorigenesis. Proceedings of the National Academy of Sciences of the United States of America 107: 9736–9741.

Lee J, Platt KA, Censullo P, Ruizi Altaba A. 1997. Gli1 is a target of Sonic hedgehog that induces ventral neural tube development. Development (Cambridge, England) 124: 2537–2552.

Lee LA, Karabina A, Broadwell LJ, Leinwand LA. 2019. The ancient sarcomeric myosins found in specialized muscles. Skelet Muscle 9: 7.

Lei Q, Zelman AK, Kuang E, Li S, Matise MP. 2004. Transduction of graded Hedgehog signaling by a combination of Gli2 and Gli3 activator functions in the developing spinal cord. Development 131: 3593–3604.

Lex RK, Ji Z, Falkenstein KN, Zhou W, Henry JL, Ji H, Vokes SA. 2020. GLI transcriptional repression regulates tissue-specific enhancer activity in response to Hedgehog signaling. Elife 9.

Li G, Jia Q, Zhao J, Li X, Yu M, Samuel MS, Zhao S, Prather RS, Li C. 2014. Dysregulation of genome-wide gene expression and DNA methylation in abnormal cloned piglets. BMC genomics 15: 811.

Li H, Durbin R. 2009. Fast and accurate short read alignment with Burrows-Wheeler transform. Bioinformatics 25: 1754–1760.

Li Q, Lex RK, Chung H, Giovanetti SM, Ji Z, Ji H, Person MD, Kim J, Vokes SA. 2016. The Pluripotency Factor NANOG Binds to GLI Proteins and Represses Hedgehog-mediated Transcription. The Journal of biological chemistry 291: 7171–7182.

Litingtung Y, Chiang C. 2000. Specification of ventral neuron types is mediated by an antagonistic interaction between Shh and Gli3. Nature neuroscience 3: 979–985.

Litingtung Y, Dahn RD, Li Y, Fallon JF, Chiang C. 2002. Shh and Gli3 are dispensable for limb skeleton formation but regulate digit number and identity. Nature 418: 979–983.

Lopez-Rios J, Duchesne A, Speziale D, Andrey G, Peterson KA, Germann P, Unal E, Liu J, Floriot S, Barbey S et al. 2014. Attenuated sensing of SHH by Ptch1 underlies evolution of bovine limbs. Nature 511: 46–51.

Lorberbaum DS, Ramos AI, Peterson KA, Carpenter BS, Parker DS, De S, Hillers LE, Blake VM, Nishi Y, McFarlane MR et al. 2016. An ancient yet flexible cis-regulatory architecture allows localized Hedgehog tuning by patched/Ptch1. Elife 5.

Marcucio RS, Tong M, Helms JA. 2001. Establishment of distinct signaling centers in the avian frontonasal process. Developmental biology 235.

Matise MP, Epstein DJ, Park HL, Platt KA, Joyner AL. 1998. Gli2 is required for induction of floor plate and adjacent cells, but not most ventral neurons in the mouse central nervous system. Development 125: 2759–2770.

Matise MP, Joyner AL. 1999. Gli genes in development and cancer. Oncogene 18: 7852–7859.

Maves L, Tyler A, Moens CB, Tapscott SJ. 2009. Pbx acts with Hand2 in early myocardial differentiation. Developmental biology 333: 409–418.

McDermott A, Gustafsson M, Elsam T, Hui CC, Emerson CP, Jr., Borycki AG. 2005. Gli2 and Gli3 have redundant and context-dependent function in skeletal muscle formation. Development 132: 345–357.

McFadden DG, McAnally J, Richardson JA, Charite J, Olson EN. 2002. Misexpression of dHAND induces ectopic digits in the developing limb bud in the absence of direct DNA binding. Development 129: 3077–3088.

Millington G, Elliott KH, Chang YT, Chang CF, Dlugosz A, Brugmann SA. 2017. Cilia-dependent GLI processing in neural crest cells is required for tongue development. Developmental biology 424: 124–137.

Mo R, Freer AM, Zinyk DL, Crackower MA, Michaud J, Heng HH, Chik KW, Shi XM, Tsui LC, Cheng SH et al. 1997. Specific and redundant functions of Gli2 and Gli3 zinc finger genes in skeletal patterning and development. *Development (Cambridge*, England) 124: 113–123.

Morikawa Y, D’Autreaux F, Gershon MD, Cserjesi P. 2007. Hand2 determines the noradrenergic phenotype in the mouse sympathetic nervous system. Developmental biology 307: 114–126.

Narasimhan K, Lambert SA, Yang AW, Riddell J, Mnaimneh S, Zheng H, Albu M, Najafabadi HS, Reece-Hoyes JS, Fuxman Bass JI et al. 2015. Mapping and analysis of Caenorhabditis elegans transcription factor sequence specificities. Elife 4.

Nusslein-Volhard C, Wieschaus E. 1980. Mutations affecting segment number and polarity in Drosophila. Nature 287: 795–801.

Olson EN, Srivastava D. 1996. Molecular pathways controlling heart development. Science 272: 671–676.

Oosterveen T, Kurdija S, Alekseenko Z, Uhde CW, Bergsland M, Sandberg M, Andersson E, Dias JM, Muhr J, Ericson J. 2012. Mechanistic differences in the transcriptional interpretation of local and long-range Shh morphogen signaling. Developmental cell 23: 1006–1019.

Osterwalder M, Speziale D, Shoukry M, Mohan R, Ivanek R, Kohler M, Beisel C, Wen X, Scales SJ, Christoffels VM et al. 2014. HAND2 targets define a network of transcriptional regulators that compartmentalize the early limb bud mesenchyme. Developmental cell 31: 345–357.

Pan Y, Bai CB, Joyner AL, Wang B. 2006. Sonic hedgehog signaling regulates Gli2 transcriptional activity by suppressing its processing and degradation. Mol Cell Biol 26: 3365–3377.

Park HL, Bai C, Platt KA, Matise MP, Beeghly A, Hui CC, Nakashima M, Joyner AL. 2000. Mouse Gli1 mutants are viable but have defects in SHH signaling in combination with a Gli2 mutation. Development 127: 1593–1605.

Parker DS, White MA, Ramos AI, Cohen BA, Barolo S. 2011. The cis-regulatory logic of Hedgehog gradient responses: key roles for gli binding affinity, competition, and cooperativity. Sci Signal 4: ra38.

Persson M, Stamataki D, te Welscher P, Andersson E, Bose J, Ruther U, Ericson J, Briscoe J. 2002. Dorsal-ventral patterning of the spinal cord requires Gli3 transcriptional repressor activity. Genes Dev 16: 2865–2878.

Peterson KA, Nishi Y, Ma W, Vedenko A, Shokri L, Zhang X, McFarlane M, Baizabal JM, Junker JP, van Oudenaarden A et al. 2012. Neural-specific Sox2 input and differential Gli-binding affinity provide context and positional information in Shh-directed neural patterning. Genes & development 26: 2802–2816.

Quinlan AR, Hall IM. 2010. BEDTools: a flexible suite of utilities for comparing genomic features. Bioinformatics 26: 841–842.

Roelink H, Augsburger A, Heemskerk J, Korzh V, Norlin S, Ruizi Altaba A. 1994. Floor plate and motor neuron induction by vhh-1, a vertebrate homolog of hedgehog ex-pressed by the notochord. Cell 76: 761–775.

Roessler E, Du YZ, Mullor JL, Casas E, Allen WP, Gillessen-Kaesbach G, Roeder ER, Ming JE, Ruizi Altaba A, Muenke M. 2003. Loss-of-function mutations in the human GLI2 gene are associated with pituitary anomalies and holoprosencephaly-like features. Proceedings of the National Academy of Sciences of the United States of America 100: 13424–13429.

Sasaki H, Hui C, Nakafuku M, Kondoh H. 1997. A binding site for Gli proteins is essential for HNF-3beta floor plate enhancer activity in transgenics and can respond to Shh in vitro. Development 124: 1313–1322.

Sasaki H, Nishizaki Y, Hui C, Nakafuku M, Kondoh H. 1999. Regulation of Gli2 and Gli3 activities by an amino-terminal repression domain: implication of Gli2 and Gli3 as primary mediators of Shh signaling. Development 126: 3915–3924.

Shimeld SM, van den Heuvel M, Dawber R, Briscoe J. 2007. An amphioxus Gli gene reveals conservation of midline patterning and the evolution of hedgehog signalling diversity in chordates. PloS one 2: e864.

Shin SH, Kogerman P, Lindstrom E, Toftgard R, Biesecker LG. 1999. GLI3 mutations in human disorders mimic Drosophila cubitus interruptus protein functions and localization. Proceedings of the National Academy of Sciences of the United States of America 96: 2880–2884.

Srivastava D, Thomas T, Lin Q, Kirby ML, Brown D, Olson EN. 1997. Regulation of cardiac mesodermal and neural crest development by the bHLH transcription factor, dHAND. Nature genetics 16: 154–160.

St-Jacques B, Hammerschmidt M, McMahon AP. 1999. Indian hedgehog signaling regulates proliferation and differentiation of chondrocytes and is essential for bone formation. Genes and Development 13: 2072–2086.

Stamataki D, Ulloa F, Tsoni SV, Mynett A, Briscoe J. 2005. A gradient of Gli activity mediates graded Sonic Hedgehog signaling in the neural tube. Genes Dev 19: 626–641.

Tan Z, Niu B, Tsang KY, Melhado IG, Ohba S, He X, Huang Y, Wang C, McMahon AP, Jauch R et al. 2018. Synergistic co-regulation and competition by a SOX9-GLI-FOXA phasic transcriptional network coordinate chondrocyte differentiation transitions. PLoS genetics 14: e1007346.

te Welscher P, Zuniga A, Kuijper S, Drenth T, Goedemans HJ, Meijlink F, Zeller R. 2002. Progression of vertebrate limb development through SHH-mediated counteraction of GLI3. Science 298: 827–830.

Thomas T, Kurihara H, Yamagishi H, Kurihara Y, Yazaki Y, Olson EN, Srivastava D. 1998. A signaling cascade involving endothelin-1, dHAND and msx1 regulates development of neural-crest-derived branchial arch mesenchyme. Development (Cambridge, England) 125: 3005–3014.

Uhl JD, Cook TA, Gebelein B. 2010. Comparing anterior and posterior Hox complex formation reveals guidelines for predicting cis-regulatory elements. Developmental biology 343: 154–166.

Uhl JD, Zandvakili A, Gebelein B. 2016. A Hox Transcription Factor Collective Binds a Highly Conserved Distal-less cis-Regulatory Module to Generate Robust Transcriptional Outcomes. PLoS genetics 12: e1005981.

Vokes SA, Ji H, McCuine S, Tenzen T, Giles S, Zhong S, Longabaugh WJ, Davidson EH, Wong WH, McMahon AP. 2007. Genomic characterization of Gli-activator targets in sonic hedgehog-mediated neural patterning. *Development (Cambridge*, England) 134: 1977–1989.

Vokes SA, Ji H, Wong WH, McMahon AP. 2008. A genome-scale analysis of the cis-regulatory circuitry underlying sonic hedgehog-mediated patterning of the mammalian limb. Genes Dev 22: 2651–2663.

Vortkamp A, Franz T, Gessler M, Grzeschik KH. 1992. Deletion of GLI3 supports the homology of the human Greig cephalopolysyndactyly syndrome (GCPS) and the mouse mutant extra toes (Xt). Mammalian Genome 3: 461–463.

Vortkamp A, Gessler M, Grzeschik KH. 1991. GLI3 zinc-finger gene interrupted by translocations in Greig syndrome families. Nature 352: 539–540.

Wang B, Fallon JF, Beachy PA. 2000. Hedgehog-regulated processing of Gli3 produces an anterior/posterior repressor gradient in the developing vertebrate limb. Cell 100: 423–434.

Wang Q, Wang G, Wang B, Yang H. 2018. Activation of TGR5 promotes osteoblastic cell differentiation and mineralization. Biomed Pharmacother 108: 1797–1803.

Wang QT, Holmgren RA. 1999. The subcellular localization and activity of Drosophila cubitus interruptus are regulated at multiple levels. Development 126: 5097–5106.

Weirauch Matthew T, Yang A, Albu M, Cote AG, Montenegro-Montero A, Drewe P, Najafabadi Hamed S, Lambert Samuel A, Mann I, Cook K et al. 2014. Determination and Inference of Eukaryotic Transcription Factor Sequence Specificity. Cell 158: 1431–1443.

Wen X, Lai CK, Evangelista M, Hongo JA, de Sauvage FJ, Scales SJ. 2010. Kinetics of hedgehog-dependent full-length Gli3 accumulation in primary cilia and subsequent degradation. Molecular and cellular biology 30: 1910–1922.

Wijgerde M, McMahon JA, Rule M, McMahon AP. 2002. A direct requirement for Hedgehog signaling for normal specification of all ventral progenitor domains in the presumptive mammalian spinal cord. Genes Dev 16: 2849–2864.

Wild A, Kalff-Suske M, Vortkamp A, Bornholdt D, Konig R, Grzeschik KH. 1997. Point mutations in human GLI3 cause Greig syndrome. Hum Mol Genet 6: 1979–1984.

Witt LM, Gutzwiller LM, Gresser AL, Li-Kroeger D, Cook TA, Gebelein B. 2010. Atonal, Senseless, and Abdominal-A regulate rhomboid enhancer activity in abdominal sensory organ precursors. Developmental biology 344: 1060–1070.

Wolpert L. 1969. Positional information and the spatial pattern of cellular differentiation. J Theor Biol 25: 1–47.

Xu J, Liu H, Lan Y, Adam M, Clouthier DE, Potter S, Jiang R. 2019. Hedgehog signaling patterns the oral-aboral axis of the mandibular arch. Elife 8.

Yao HH, Whoriskey W, Capel B. 2002. Desert Hedgehog/Patched 1 signaling specifies fetal Leydig cell fate in testis organogenesis. Genes Dev 16: 1433–1440.

Yelon D, Ticho B, Halpern ME, Ruvinsky I, Ho RK, Silver LM, Stainier DY. 2000. The bHLH transcription factor hand2 plays parallel roles in zebrafish heart and pectoral fin development. Development 127: 2573–2582.

Yin WC, Satkunendran T, Mo R, Morrissy S, Zhang X, Huang ES, Uuskula-Reimand L, Hou H, Son JE, Liu W et al. 2019. Dual Regulatory Functions of SUFU and Targetome of GLI2 in SHH Subgroup Medulloblastoma. Developmental cell 48: 167–183 e165.

Young NM, Chong HJ, Hu D, Hallgrimsson B, Marcucio RS. 2010. Quantitative analyses link modulation of sonic hedgehog signaling to continuous variation in facial growth and shape. Development 137: 3405–3409.

Zhang L, Chen XM, Sun ZJ, Bian Z, Fan MW, Chen Z. 2006. Epithelial expression of SHH signaling pathway in odontogenic tumors. Oral Oncol 42: 398–408.

Zhang X, McGrath PS, Salomone J, Rahal M, McCauley HA, Schweitzer J, Kovall R, Gebelein B, Wells JM. 2019. A Comprehensive Structure-Function Study of Neurogenin3 Disease-Causing Alleles during Human Pancreas and Intestinal Organoid Development. Dev Cell 50: 367–380 e367.

Zhu L, Zhou G, Poole S, Belmont JW. 2008. Characterization of the interactions of human ZIC3 mutants with GLI3. Human mutation 29: 99–105.

